# The Association Between Poor Sleep and Accelerated Brain Ageing

**DOI:** 10.1101/2021.06.16.448332

**Authors:** Jivesh Ramduny, Matteo Bastiani, Robin Huedepohl, Stamatios N. Sotiropoulos, Magdalena Chechlacz

**Affiliations:** Sir Peter Mansfield Imaging Centre, School of Medicine, University of Nottingham, Nottingham, UK; School of Psychology, Trinity College Dublin, Dublin, Ireland; Trinity College Institute of Neuroscience, Trinity College Dublin, Dublin, Ireland; National Institute for Health Research (NIHR), Nottingham Biomedical Research Centre, Queen’s Medical Centre, Nottingham, UK; Wellcome Centre for Integrative Neuroimaging (WIN) – Oxford Centre for Functional Magnetic Resonance Imaging of the Brain (FMRIB), University of Oxford, Oxford, UK; School of Psychology, University of Birmingham, Birmingham, UK; Centre for Human Brain Health, University of Birmingham, Birmingham, UK

**Author notes:** Equal contribution (Stamatios N. Sotiropoulos, Magdalena Chechlacz).

**Keywords:** sleep, ageing, brain age, gray matter, white matter, magnetic resonance imaging

## Abstract

The ageing brain undergoes widespread gray (GM) and white matter (WM) degeneration. But numerous studies indicate large heterogeneity in the age-related brain changes, which can be attributed to modifiable lifestyle factors, including sleep. Inadequate sleep has been previously linked to GM atrophy and WM changes. However, the reported findings are highly inconsistent. By contrast to previous research independently characterizing patterns of either the GM or the WM changes, we used here linked independent component analysis (FLICA) to examine covariation in GM and WM in a group of older adults. Next, we employed a novel technique to estimate the brain age delta (i.e. difference between chronological and apparent brain age assessed using neuroimaging data) and study its associations with sleep quality and sleep fragmentation, hypothesizing that poor sleep accelerates brain ageing. FLICA revealed a number of multimodal (including both GM and WM) neuroimaging components, associated with age, but also with sleep quality and sleep fragmentation. Brain age delta estimates were highly sensitive in detecting the effects of sleep problems on the ageing brain. Specifically, we show significant associations between brain age delta and poor sleep quality, suggesting two years deviation above the chronological age. Our findings indicate that sleep problems in healthy older adults should be considered a risk factor for accelerated brain ageing.

## 1 INTRODUCTION

The ageing brain undergoes widespread gray (GM) and white matter (WM) degeneration associated with functional brain changes and gradual cognitive decline. Numerous studies indicate large heterogeneity in the age-related brain changes in older adults (Cabeza et al., 2017; Eavani et al., 2018; Good et al., 2001; Raz & Rodrigue, 2006; Westlye et al., 2010). Understanding this heterogeneity is of high importance as it could provide key insights into why some older adults go through rapid cognitive deterioration progressing to dementia, while others experience only mild decline in cognitive functioning or no noticeable cognitive changes (Hayden et al., 2011). However, disentangling the heterogeneity in brain ageing has proved to be a challenge despite numerous research efforts. Some of the ongoing work focuses on phenotyping to capture a variation in brain ageing and predict trajectories of normal versus pathological ageing (e.g., Eavani et al., 2018). Other studies explore compensatory and neuroadaptive mechanisms often linked to lifelong cumulative cognitive engagement (for review see Cabeza et al., 2018; Fabiani, 2012; Stern et al., 2020). Finally, a growing body of work links heterogeneity in brain ageing to modifiable lifestyle factors such as sleep, diet and physical activity (Wassenaar et al., 2019).

Data analysis techniques rooted in structural magnetic resonance imaging (MRI), such as voxel-based morphometry (Ashburner & Friston, 2000) and cortical surface analysis (Dale et al., 1999; Fischl, 2012; Fischl & Dale, 2000) from T1-weighted scans, have been fundamental in identifying volumetric gray matter and cortical thickness changes, respectively, in relation to age-related cognitive decline (Giorgio et al., 2010; Good et al., 2001; Lemaitre et al., 2012). Additionally, diffusion MRI (dMRI)-based measures sensitive to microstructural properties of white matter have been successfully employed to characterize white matter alterations and changes in structural connectivity in relation to cognitive ageing (e.g., Barrick et al., 2010; Burzynska et al., 2010; Giorgio et al., 2010). Such approaches characterize patterns of either gray or white matter changes independently, estimated from a single MRI modality. A potential shortcoming is that they fail to capture interlinked gray and white matter changes associated with cognitive decline as well as to model covariation in gray and white matter age-related deterioration. Furthermore, information provided by methods based on different modalities might be difficult to integrate into a single model of brain ageing and occasionally the resulting findings seem contradictory (e.g., McGinnis et al., 2011; Raz & Rodrigue, 2006). To overcome these drawbacks and limitations methods aimed at fusing information from multiple modalities have been developed (e.g., Groves et al., 2011; Liu et al., 2009; Xu et al., 2009). One of such methods, based on linked independent component analysis (Groves et al., 2011), enables decomposition of data from multiple modalities into spatial components to model variation in cross-modal imaging features across groups of participants. Linked independent component analysis has been previously applied to access patterns of structural brain changes during healthy and pathological ageing (Douaud et al., 2014; Groves et al., 2011; Groves et al., 2012).

Another approach to capturing the inter-individual differences in the rate of brain ageing is based on estimation of so called “brain age gap” or “brain age delta” i.e., difference between “chronological age” calculated from the date of birth and “brain age” computed based on neuroimaging data (Cole & Franke, 2017; Cole et al., 2017; Franke & Gaser, 2019; Liem et al., 2017; Smith et al., 2019). As this method enables to quantify the deviation from normative ageing, it has been used as a biomarker of brain ageing; to assess accelerated brain ageing associated with Alzheimer’s disease as well as a predictor of progression from mild cognitive impairment to dementia (Boyle et al., 2021; Cole & Franke, 2017; Franke & Gaser, 2019; Gaser et al., 2013). Similarly, this method has a potential to provide understanding of long term predictors of brain health, including socio-demographic and lifestyle factors. One recent study examined the effect of education and physical activity on the gap between “chronological age” and “brain age”, elegantly demonstrating that higher levels of education and physical activity have a positive impact on the ageing brain, supporting more “youthful” state (Steffener et al., 2016).

Sleep disruptions constitute a potentially modifiable risk factor for dementia and reduced longevity. As we get older overall sleep quality deteriorates. Up to half of elderly population experiences various sleep problems and sleep disruptions, including difficulties in maintaining or initiating sleep, and fragmentation of sleep (Miner & Kryger, 2017; Varma et al., 2019). Short sleep duration and poor sleep quality in healthy older adults have been previously associated with grey matter atrophy and microstructural white matter changes, consequently leading to cognitive decline (Sexton et al., 2014; Sexton et al., 2017; Wassenaar et al., 2019). However, the reports linking sleep problems, brain changes, cognitive outcomes and overall increased risk of dementia are inconsistent (Sexton et al., 2020; Zitser et al., 2020). For example, a recent longitudinal ageing study, spanning over almost three decades failed to demonstrate a significant link between sleep duration and either grey matter or white matter microstructure (Zitser et al., 2020). To our best knowledge all the previous studies separately examined patterns of either gray or white matter changes estimated from a single MRI modality.

In this study, using linked independent component analysis, we explored interconnected GM and WM microstructural changes due to brain aging and sleep problems in a group of 50 healthy elderly participants using measures extracted from structural and diffusion MRI. We investigated the associations of age-related brain changes with two measures indicative of sleep problems, sleep quality index and sleep fragmentation, hypothesizing that poor sleep accelerates brain ageing. We demonstrate that linked analysis of multimodal imaging data reveals associations between brain structural and microstructural features and poor sleep indicators, which are not evident using conventional unimodal analysis. Furthermore, to assess accelerated brain ageing, we employed a recently-introduced technique (Smith et al., 2019) to estimate brain age delta, the deviation from chronological brain age, in an unbiased manner (i.e., by applying linear and quadratic correction to remove age-related biases). While the most widely used approaches to study “brain age gap” (for review see Cole & Franke, 2017; Franke & Gaser, 2019) are based on predictors derived from a single MRI modality, we employed a multimodal approach, with a set of structural and microstructural imaging-derived features (Smith et al., 2019). We show significant associations between this unbiased brain age delta and poor sleep quality, suggesting two years deviation above the chronological age for poor sleepers.

## 2 METHODS AND MATERIALS

### 2.1 Participants

Fifty older adults participated in the study (22 males; age range 65-84; mean ± SD age 73.5 ± 4.7). All participants were recruited either from the Neuropsychological panel of elderly volunteers, or the Birmingham 1000 Elders group, both established at the University of Birmingham. The two panels of elderly volunteers consist of adults aged 65 or over who are in good health and have no pre-existing cognitive impairment. All participants had normal or corrected-to-normal vision, had no history of psychiatric or neurological disease and were right-handed (self-report). Participants with contraindications to MRI were excluded. The study was approved by the University of Birmingham Ethical Review Committee and all participants provided written informed consent and received monetary compensation for participation in agreement with approved ethics protocol. The demographic characteristics of participants are presented in Table 1.

**Table 1.** Participants characteristics

| Variable | Number (N=50) | Mean (SD) | Range |
| --- | --- | --- | --- |
| Age (years) | — | 73.5 (4.7) | 65-84 |
| 65-69 | 11 | — | — |
| 70-74 | 17 | — | — |
| 75-79 | 16 | — | — |
| 80-85 | 6 | — | — |
| Gender (Male/Female) | 22/28 | — | — |
| PSQI (global score)‡ | — | 5.6 (3.3) | 0-15 |
| Subjective sleep quality* | — | 0.9 (0.6) | 0-2 |
| Sleep latency* | — | 0.9 (0.9) | 0-3 |
| Sleep duration* | — | 0.8 (0.7) | 0-2 |
| Sleep efficiency* | — | 0.9 (0.9) | 0-3 |
| Sleep disruptions* | — | 1.3 (0.5) | 0-2 |
| Sleep medication* | — | 0.3 (0.6) | 0-3 |
| Daytime dysfunctions* | — | 0.8 (0.6) | 0-3 |
| WASO (minutes) | — | 39.6 (19.2) | 15-107 |

PSQI = Pittsburgh Sleep Quality Index (‡ maximum score 21; \*maximum score 3); WASO = wake after sleep onset.

### 2.2 Sleep Assessment

Wrist actigraphy, sleep diaries and the Pittsburgh Sleep Quality Index (PSQI) questionnaire were used as objective and self-reported/subjective measures of habitual sleep. To objectively evaluate sleep patterns, participants were asked to wear wrist actigraphs (Actiwatch2, Philips Respironics Ltd) for a period of two weeks prior to the scheduled MRI scanning session. Actigraphs were set to one minute epochs (medium sensitivity setting), and collected data analyzed using Respironics Actiware 6 (Philips, Netherlands) software, which translates wrist movement data into sleep scores (sleep/wake cycles). The software output was validated using sleep diaries completed alongside actigraphy. For the purpose of the current study, we used Actiware outputs to calculate wake after sleep onset (WASO), a parameter measuring wakefulness time (in minutes) occurring after sleep onset, which indexes sleep fragmentation. In addition, participants were asked to self-evaluate their sleep quality by completing the PSQI questionnaire (Buysse et al., 1989). PSQI is a validated scale consisting of 19 self-rated questions assessing sleep quality and sleep problems over a period of one month. Each question is rated on a 4-point Likert scale from 0 (not during the past month) to 3 (three or more times a week). The scores are first combined into seven components (subjective sleep quality, sleep latency, sleep duration, habitual sleep efficiency, sleep disturbances, use of sleeping medication and daytime dysfunction), which are added to produce a global score of sleep quality, with a range of 0-21 points, with a higher score indexing worse sleep quality (Buysse et al., 1989; Carpenter & Andrykowski, 1998). PSQI has been shown to have a good reliability and validity in assessing sleep quality in the elderly population (Buysse et al., 1991; Gentili et al., 1995). In order to ensure normal distribution, the WASO and PSQI global scores were log and square root transformed, respectively, prior to using them in the subsequent statistical analyses as measures of sleep fragmentation and sleep quality.

### 2.3 MRI data acquisition

Structural T1-weighted and diffusion-weighted scans were acquired at the Birmingham University Imaging Centre (BUIC) using a Philips 3T Achieva scanner a 32-channel head coil. A T1-weighted MPRAGE with spatial resolution 1x1x1mm^3^ (176 sagittal slices, TR = 7.5 ms, TE = 3.5 ms, flip angle = 8°) was obtained for each participant, along with a multi-shell dMRI (single-shot EPI, 2×2×2mm^3^, TR= 9000 ms, TE= 81.5 ms, 5 x b=0 s/mm^2^, 50 x b=1000s/mm^2^, 50 x b=2000s/mm^2^, plus 5 x b=0 s/mm^2^ phase encoding-reversed to correct for susceptibility-induced artifacts; (Andersson et al., 2003).

### 2.4 T1-weighted data pre-processing and Voxel-Based Morphometry

The structural T1-weighted images were preprocessed using the UK Biobank T1-weighted pipeline (Alfaro-Almagro et al., 2018). The pipeline corrects for bias fields, performs skull-stripping and aligns data to the MNI152 standard template, before segmenting the T1 images into different tissue classes (e.g. GM/WM/CSF) as well as into to cortical and subcortical structures.

To examine whole brain effects of ageing and poor sleep (sleep problems) on GM structure, the brain-extracted images were processed following the FSL voxel-based morphometry (VBM) pipeline (Douaud et al., 2007; Good et al., 2001). The resulting GM images were first non-linearly registered to the MNI152 standard space and they were concatenated and averaged to create a study-specific GM template. All of the 50 native images were then non-linearly registered to this study-specific template and they were modulated by the determinant of the Jacobian of the non-linear warp field to correct for local enlargement or contraction due to the transformation. The 50 modulated registered GM images were smoothed with an isotropic Gaussian kernel with a sigma of 2mm (~5mm FWHM). We examined widespread GM volumetric changes in the brain by building a general linear model (GLM) using the demeaned age as a regressor. We also studied the effects of sleep problems on the GM changes by building two GLMs using the demeaned PSQI and WASO while regressing out age, respectively. We used FSL’s randomise to perform non-parametric inference (Winkler et al., 2014). Threshold-free cluster enhancement (TFCE) was applied to avoid the selection of arbitrary initial cluster-forming threshold (Smith & Nichols, 2009) and 1000 permutations were performed for each contrast defined in the GLMs. The statistical maps were corrected for multiple comparisons using family-wise error rate (FWE).

### 2.5 Extraction of T1 features

We extracted 110 structural (T1) imaging derived phenotypes (T1 IDPs) using the UK Biobank T1 pipeline (Alfaro-Almagro et al., 2018). The T1 IDPs represented volumes of cortical and subcortical structures in each hemisphere in the standard MNI152 space which were based on the Harvard-Oxford structural atlases (https://fsl.fmrib.ox.ac.uk/fsl/fslwiki/Atlases). The cortical IDPs were obtained using FAST (Zhang et al., 2001) to derive the GM volumes of the cortical regions of interest (ROIs). The subcortical IDPs were obtained using FMRIB’s Integrated Registration and Segmentation Tool (FIRST; Patenaude et al., 2011) to produce the GM volumes of the subcortical ROIs including the limbic, basal ganglia and thalamic sub-regions, and extending to the brainstem. These T1-derived features, along with the dMRI-derived features (see section 2.7), were used for brain age delta estimation (section 2.9). The complete list of the 110 T1 IDPs that were used in the brain age delta models is presented in Supplementary Table 1.

### 2.6 Diffusion data pre-processing and Tract-based Spatial Statistics

The dMRI data were preprocessed using the UK biobank pipeline (Alfaro-Almagro et al., 2018). The procedure corrects for susceptibility induced distortion, eddy-current distortion and motion using the EDDY toolbox (Andersson & Sotiropoulos, 2016) and obtains transformations of the diffusion to structural and standard space. A diffusion tensor model (Basser et al., 1994) was fitted to low b-value (b=0 and 1000s/mm^2^) shells of each voxel of the corrected diffusion data to obtain microstructural maps including fractional anisotropy (FA) and mean diffusivity (MD).

To examine whole brain effects of ageing and sleep problems on WM microstructure, Tract-Based Spatial Statistics (TBSS) were carried out to skeletonize and transform the FA volumes into a common space (Smith et al., 2006). The FA native images were non-linearly registered to the FMRIB58 FA standard space using FNIRT due to its good native-to-standard warping across different age groups (Andersson et al., 2019; Westlye et al., 2010). The mean FA volume from the 50 subjects was derived and thinned to create a study-specific mean FA skeleton which represents the centers of all common tracts. The mean FA skeleton was then thresholded and binarised at FA > 0.2 to minimize partial volume effects with the boundaries of GM and CSF tissues. The subject-wise FA volumes were warped onto this mean FA skeleton to produce skeletonized FA data and the same warping procedure was applied to the MD maps to yield skeletonized MD data from voxels with FA > 0.2. The resulting skeletonized FA and MD maps were then fed into voxelwise cross-subject statistics. We constructed GLMs to test for widespread effects of ageing on FA and MD while regressing out the effects of motion, as estimated by EDDY. We also examined the effects of poor sleep on FA and MD by building GLMs using the demeaned PSQI and WASO while regressing out motion and age. Threshold-free cluster enhancement (TFCE) was applied to avoid the selection of arbitrary initial cluster-forming threshold (Smith & Nichols, 2009) and 1000 permutations were performed for each contrast defined in the GLMs. The statistical maps were FWE-corrected for multiple comparisons.

### 2.7 Extraction of dMRI features

We performed automated probabilistic tractography using predefined protocols for identifying major WM tracts in the left and right hemispheres as described in FSL’s XTRACT tool (Warrington et al., 2020; https://fsl.fmrib.ox.ac.uk/fsl/fslwiki/XTRACT). Prior to XTRACT, we fitted the crossing fibre model (FSL’s BEDPOSTX; Behrens et al., 2007) to each subject’s data to estimate up to three fibre orientations per voxel. XTRACT was then used to reconstruct a set of tracts including projection, association, commissural and limbic fibre bundles for each subject. The tract probability density maps, normalized by the total number of valid streamlines, were thresholded at 0.1% and binarised to produce a tract mask for each tract in standard space. We then applied the tract mask of each subject with the TBSS derived FA skeleton to produce subject-wise skeletonised tract masks, depicting the core of each WM bundle. For each subject, the mean FA and MD within each skeletonised tract were obtained to produce 66 (33 FA and 33 MD) subject-wise microstructural IDPs. The predefined list of tracts that were reconstructed using XTRACT was: anterior commissure (AC), bilateral arcuate fasciculus (AF), bilateral acoustic radiation (AR), bilateral anterior thalamic radiation (ATR), bilateral cingulum (Cing), bilateral frontal aslant (FA), forceps major (FMA), forceps minor (FMI), fornix (FX), bilateral inferior fronto-occipital fasciculus (IFOF), bilateral inferior longitudinal fasciculus (ILF), middle cerebellar peduncle (MCP), bilateral middle longitudinal fasciculus (MdLF), bilateral optic radiation (OR), bilateral superior longitudinal fasciculus 1 (SLF1), bilateral superior longitudinal fasciculus 2 (SLF2), bilateral superior longitudinal fasciculus 3 (SLF3), bilateral superior thalamic radiation (STR) and bilateral uncinate fasciculus (UF).

### 2.8 Linked Independent Component Analysis

To jointly explore whole brain effects of ageing and sleep quality on GM structure and WM microstructure, we used the VBM and TBSS maps, extracted from T1 and diffusion MRI data respectively, as inputs to linked Independent Component Analysis (FLICA). FLICA is a data-driven approach which automatically decomposes multimodal data into independent components (ICs; Groves et al., 2011). Each IC characterizes a mode of inter-subject multimodal (e.g. brain GM structure and WM microstructure in our case) variability such that each subject loading, which is shared across the modalities, corresponds to statistically independent and non-Gaussian multi- modal spatial maps (Groves et al., 2012). Importantly, an IC’s subject loading may be dominated by a single modality as opposed to equal contribution of the modalities of interest. Given the sample size restriction, we ran a FLICA decomposition with 10 ICs on 3 inputs, derived from VBM (modulated GM density maps for each subject using a study-specific template in MNI152 space) and TBSS (skeletonised FA map and skeletonised MD map for each subject in MNI152 space) data to identify post-hoc ICs which may linearly and/or quadratically associate with age, PSQI and WASO. We evaluated statistical significance as well as plausible relationships between each IC’s loading and non-imaging measures using effect magnitude (R^2^) in addition to corrected p values (Douaud et al., 2014). The results were Bonferroni corrected for multiple comparisons across all 10 ICs.

### 2.9 Computation of Brain Age Delta

Using multimodal features extracted from GM and WM, we estimated brain age delta (δ) by employing multiple regression models as recently described in Smith et al (2019). This method allows for a linear and quadratic correction of δ, thus ensuring complete independence between δ and chronological age. For a vector Y (Nx1) representing chronological age for N subjects, we created an imaging matrix X (NxM) which denotes the summary measures of the structural and microstructural features in the studied group of elderly participants (N = 50). We computed δ by using: a) unimodal (i.e., 110 T1 or 66 dMRI) IDPs and b) multimodal (i.e., 176 T1+dMRI) IDPs, respectively (Figure 1). The head size scaling factor was derived as an additional structural IDP in accordance with the UK Biobank T1 pipeline (Alfaro-Almagro et al., 2018) and introduced as a confound (Miller et al., 2016). Next, the imaging feature matrix X was dimensionality reduced by applying singular value decomposition (SVD). We then performed 10-fold cross validation to prevent the regression models from overfitting the imaging data and estimated δ, in a manner that makes it orthogonal to age and bias-free (Smith et al., 2019). Finally, to account for potential accelerated effects of ageing on imaging measures (advanced ageing), we estimated the quadratic correction of delta by adding a non-linear (quadratic) term in the multiple regression models. As a result, δ_b_ was used to denote the biased (uncorrected) brain age delta (typically used in previous studies), δ was used to represent the linearly-corrected estimate of brain age delta, and finally, δ_q_ was used to represent the quadratically-corrected estimate of brain age delta. All models are shown in Figure 1.

**Figure 1.**
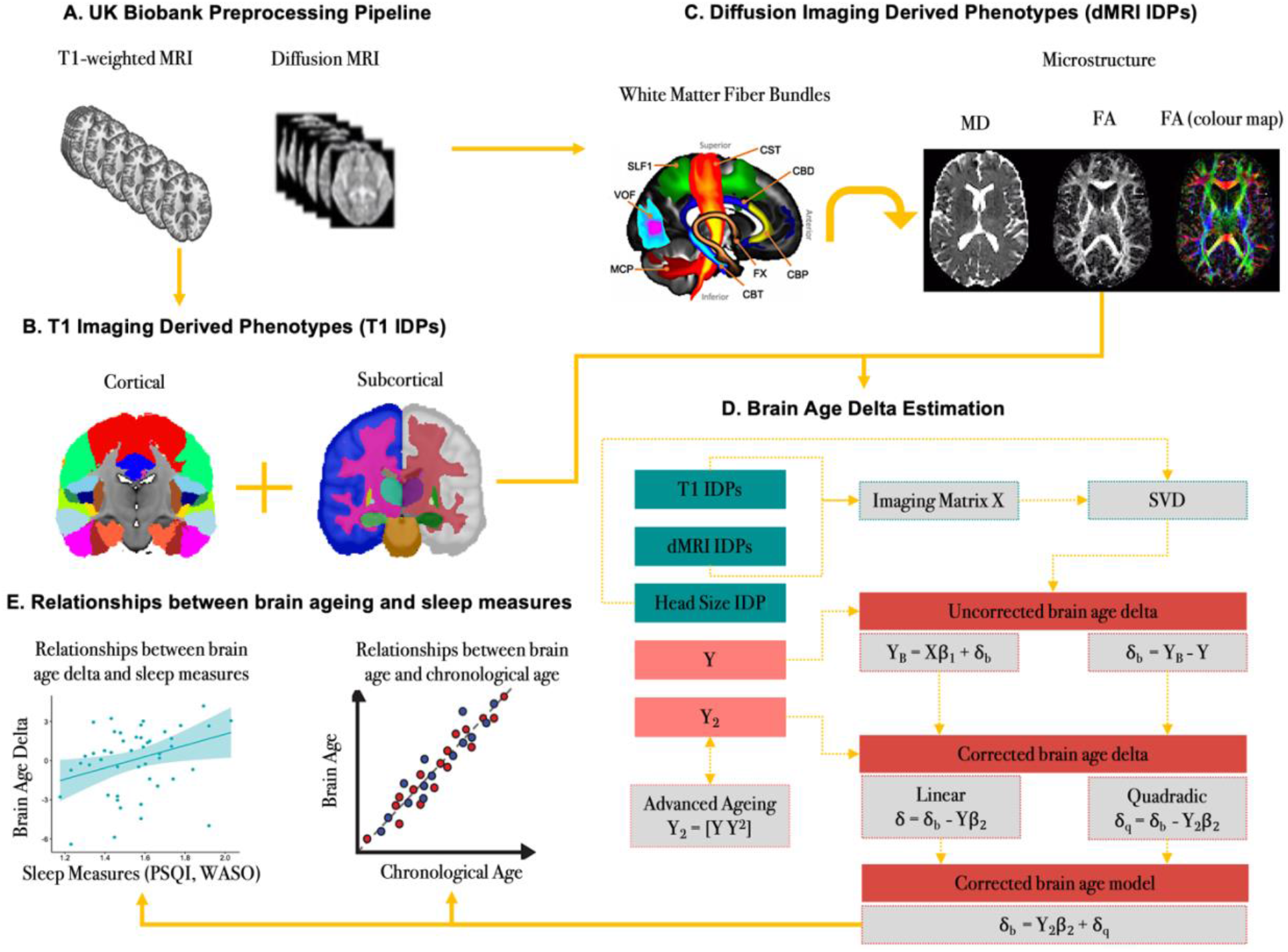
Overview of the brain age prediction model. A recently published approach described by Smith et al. (2019) was used to estimate brain age delta in an unbiased manner. **(A)** The structural and diffusion MRI data were preprocessed using the UK Biobank pipeline. **(B)** The T1 imaging derived phenotypes (T1IDPs) were extracted from the parcellations of cortical and subcortical GM volumes based on the Harvard-Oxford structural atlases. **(C)** The diffusion imaging derived phenotypes (dMRI IDPs) were extracted using predefined protocols for identifying major WM tracts as described by the FSL’s XTRACT tool. **(D)** The structural and microstructural IDPs were represented using an imaging matrix X (NxM) such that N=50 subjects and M=176 (demeaned) imaging features. The head scaling factor was used as an additional structural IDP and introduced as a confound variable in the brain age prediction model. Next, a matrix Y was created to represent the (demeaned) chronological age for N = 50 subjects. A matrix Y2 was also computed to account for quadratic ageing processes which was subsequently demeaned and orthogonalised with respect to Y. The initial brain age prediction model is Y_B_ = Xβ_1_ + δ_b_ such that β_1_ = X^+^Y and the uncorrected (biased) delta which is typically used in brain ageing studies is δ_b_ = Y_B_ – Y. The corrected (unbiased) brain age prediction model is then computed as follows: δ_b_ = Y_2_β_2_ + δ_q_ such that β_2_ = Y_2_^+^ δ_b_. The linearly-corrected estimate of delta is δ = δ_b_ – Yβ_2_ and the quadratically-corrected estimate of delta is δ_q_ = δ_b_ – Y_2_β_2_. **(E)** The relationships between chronological age (Y) and brain age (uncorrected [Y_B_]; linear [Y_L_]; quadradic [Y_Q_]) were assessed using Pearson’s correlation coefficient. The relationships between brain age delta (uncorrected [δ_b_], linear [δ], quadratic [δ_q_]) and sleep measures (PSQI, WASO) were also evaluated using Pearson’s correlation coefficient.

The R statistical package (http://r-pkgs.had.co.nz/intro.html) was used for statistical analyses of the brain age delta models with sleep patterns (i.e., PSQI, WASO) (e.g. obtaining R^2^). Similarly, we assessed the correlation strength of chronological age (Y) with the uncorrected brain age (Y_B_), the predicted linear (Y_L_) and quadratic (Y_Q_) brain age estimates. To correct for multiple comparisons, an False Discovery Rate (FDR)-corrected threshold of p < 0.05 was applied.

## RESULTS

### 3.1 VBM: GM volumetric alterations in the ageing brain

First, unimodal VBM and TBSS analyses were run to confirm that our data demonstrate anticipated trends in GM structure and WM microstructure with increasing age, as reported in previous studies. VBM analyses showed significant widespread GM volume reductions in our ageing cohort by recruiting the default mode network, primary motor cortex, primary somatosensory cortex, auditory cortex in addition to several language and visual related areas (p < 0.05 corrected; Figure 2). Significant morphological reductions were also associated with increasing age in several prefrontal, limbic and cerebellar structures (p < 0.05 corrected; Figure 2). In agreement with the literature, similar GM alterations were observed in the basal ganglia which extended to the thalamus (p < 0.05 corrected; Figure 2).

**Figure 2.**
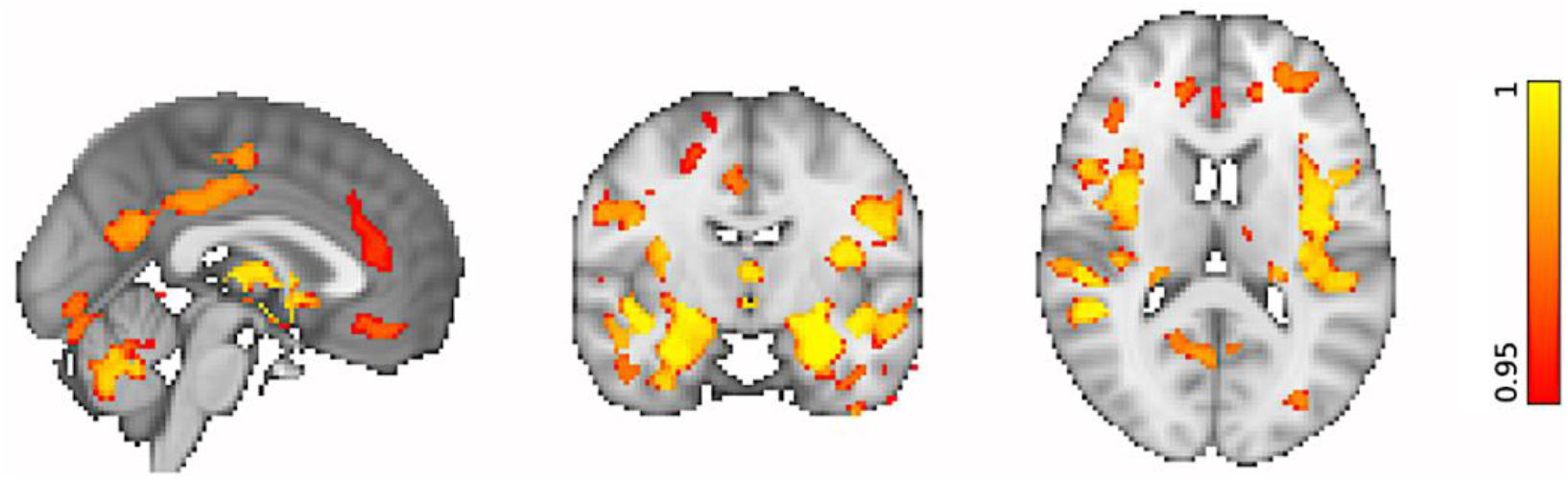
VBM: Whole-brain morphological alterations in the ageing brain. Reduced GM volumes are significantly associated with increasing age. The coloured voxels show widespread GM changes in the following regions: *Default mode network*: posterior cingulate cortex (PCC), precuneus (Pre), parahippocampal gyrus (PHG); *Primary motor cortex:* precentral gyrus (PreG); *Primary somatosensory cortex:* postcentral gyrus (PosG); *Auditory cortex:* Heschl’s gyri (HG), superior temporal gyrus (STG), planum polare (PP); *Visual areas:* lingual gyrus (LG), fusiform gyrus (FG), inferior temporal gyrus (ITG); *Language areas:* angular gyrus (AG), inferior frontal gyrus (IFG), insular cortex (INC), middle temporal gyrus (MTG); *Prefrontal regions:* superior frontal gyrus (SFG), middle frontal gyrus (MFG), frontal polare (FP); *Limbic regions:* hippocampus (HIP), amygdala (AMYG), orbitofrontal cortex (OFC); *Basal ganglia:* dorsal striatum (caudate (CAU), putamen (PUT), ventral striatum (nucleus accumbens (NAcc)), pallidum (Pa). The spatial map is corrected for multiple comparisons set at p < 0.05.

### 3.2 VBM: Sleep-related effects on GM volumes in the ageing brain

When we examined the effects of sleep quality as indexed by PSQI on whole-brain morphometry, there was no evidence of significant associations with the GM volumes, after correcting for multiple comparisons. Similarly, no associations could be found when we tested the effects of sleep fragmentation as measured by WASO on the GM volumes. Taken together, the unimodal VBM analyses did not reveal sleep problems-related alterations in whole-brain morphometry in the ageing brain.

### 3.3 TBSS: Age-related microstructural brain changes in WM

We then explored whether our cohort data support previous findings for widespread microstructural changes due to ageing. TBSS analyses demonstrated widespread age-related changes in the WM microstructure. Increasing age was significantly associated with reduced FA and increased MD (p < 0.05 corrected) within the association, projection, limbic and commissural WM fibre bundles in the studied group of neurotypical (healthy) older adults as illustrated in Figure 3.

**Figure 3.**
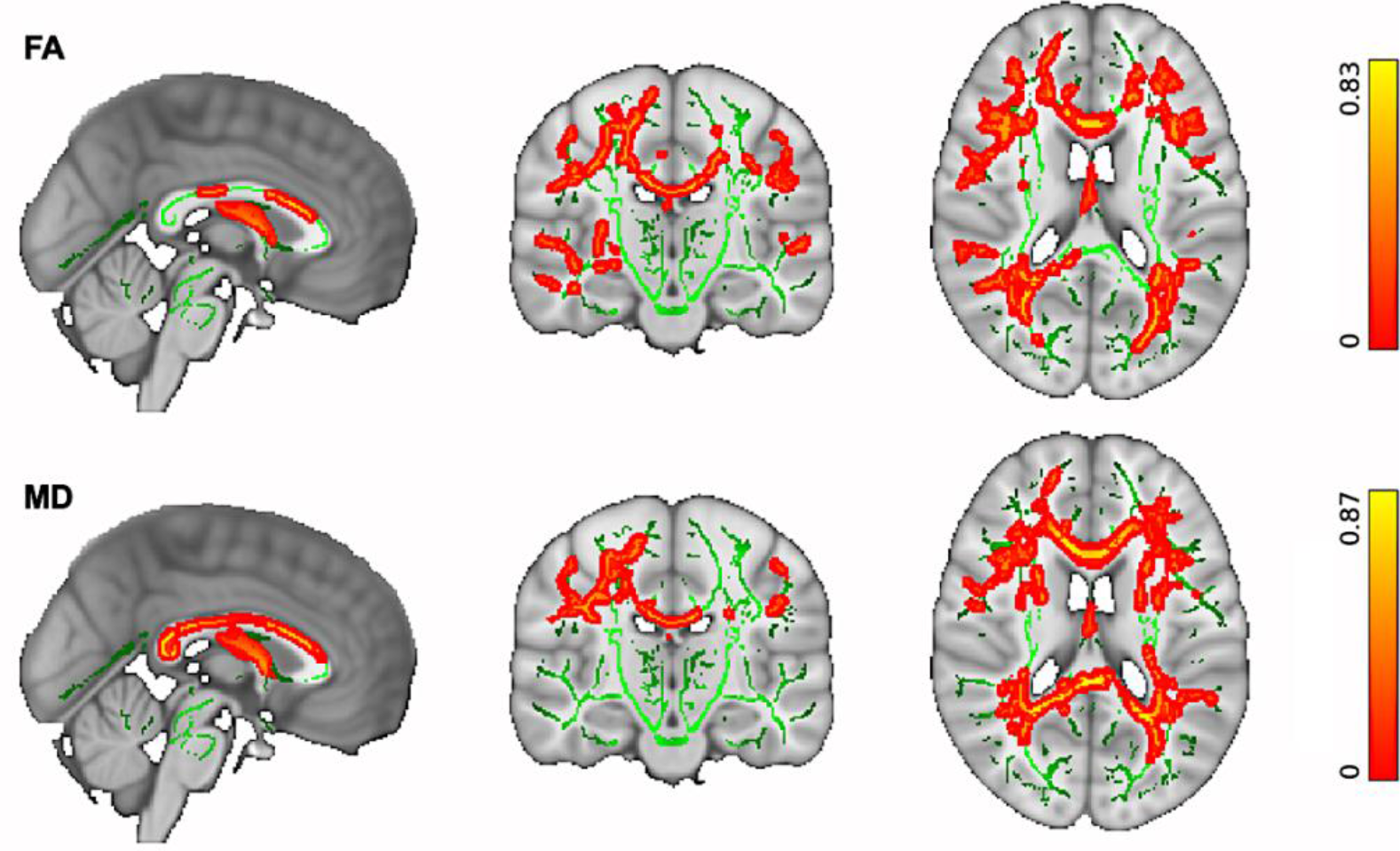
TBSS: Whole-brain skeletonised microstructural differences in older adults. (Top) Reduced FA is significantly associated with increasing age. (Bottom) Increased MD is significantly associated with increasing age in older adults. Changes were significant in a number of WM fibre bundles, including: *Association fibre bundles:* Inferior Longitudinal Fasciculus (ILF), Inferior Fronto-Occipital Fasciculus (IFOF), Superior Longitudinal Fasciculus (SLF), Uncinate Fasciculus (UF). *Projection fibre bundles:* Acoustic Radiation (AR), Anterior Thalamic Radiation (ATR), Posterior Thalamic Radiation (PTR), Corticospinal Tract (CST), Optic Radiation (OR). *Limbic fibre bundles:* Cingulum Gyrus part of Cingulum (Cing), Hippocampal part of Cingulum (CBH), Fornix (FX). *Commissural fibre bundles:* Forceps Major (FMA), Forceps Minor (FMI). The spatial maps are FDR corrected for multiple comparisons set at p < 0.05.

### 3.4 TBSS: Sleep-related effects on WM microstructure in the ageing brain

When we performed the unimodal TBSS analyses to examine the effects of sleep quality on FA or MD, there was no association between WM microstructure changes in the ageing brain with PSQI, after applying multiple comparison corrections. Similarly, no association was found when we investigated the effects of sleep fragmentation on FA and MD. Overall, the unimodal TBSS analyses did not reveal sleep problems-related changes in WM microstructure in older adults.

### 3.5 FLICA: Covariation in GM and WM changes associated with age

Similarly to the unimodal TBSS and VBM analyses, multimodal FLICA analysis revealed associations between changes in GM structure and WM microstructure and age. Two components (i.e., IC_1_ and IC_2_) were significantly associated with increasing age. IC_1_ exhibited a U-shape profile with increasing age which was related to regional GM reduction and WM changes, i.e., FA decrease and MD increase (Figure 4A). The IC_1_ subject loadings displayed a nonlinear pattern which decreased from 65 to 73 years and then increased from 73 to 84 years. While age explained 32% of the variance within IC_1_ (R^2^ = 0.32, p < 0.001 corrected), the U-shape relationship between the IC_1_ subject loading and age was dominated by MD (49%) and FA (34%) followed by GM volume (15%). The second age-related component, IC_2_, showed a linear decrease of global GM and WM microstructure with increasing age (Figure 4B). Age explained 40% of the variance within IC_2_ (R^2^ = 0.40, p < 0.001 corrected). The linear relationship between the IC_2_ loading and age was mainly driven by the GM volume (42%) followed by FA (35%) and MD (22%).

**Figure 4.**
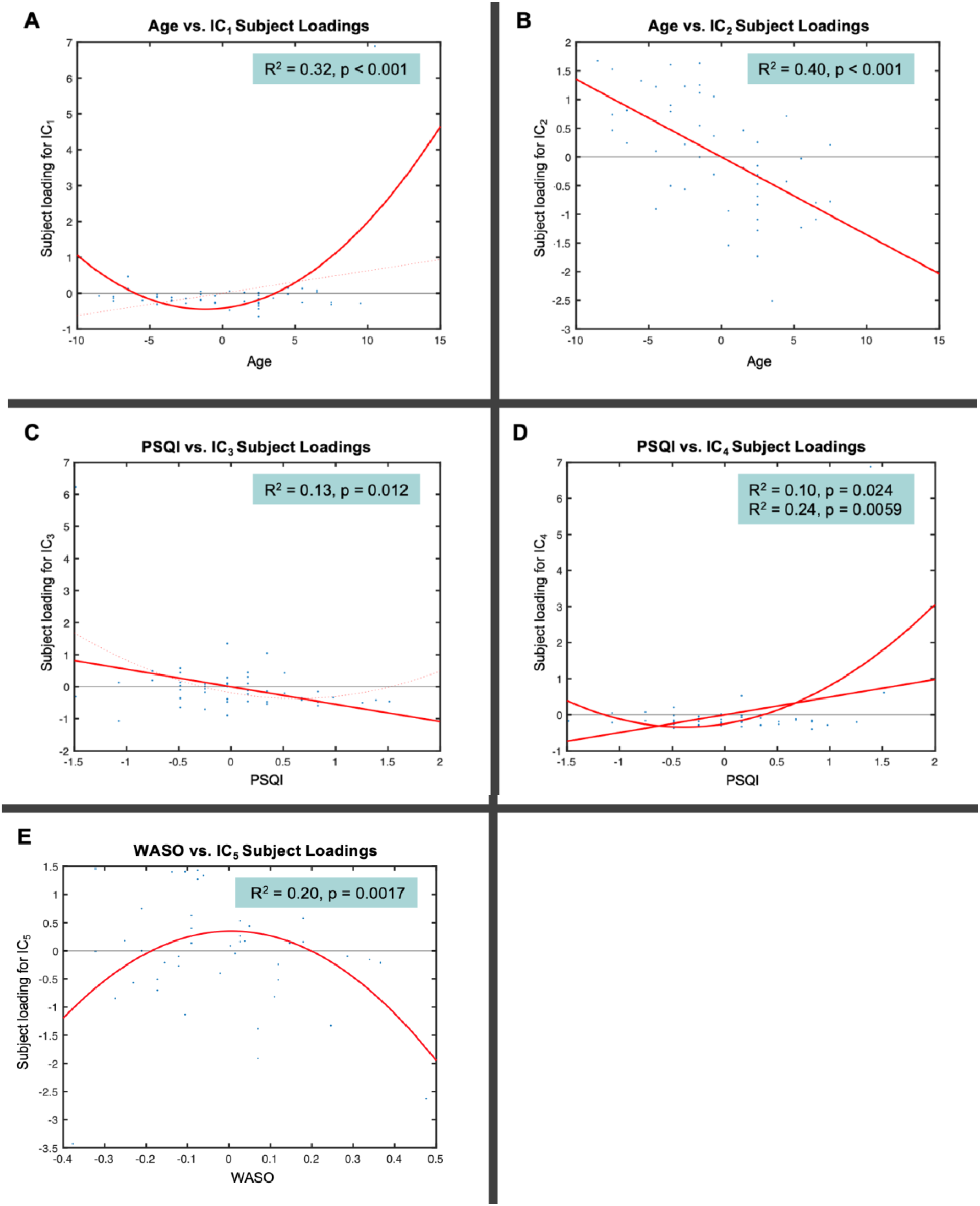
FLICA: Strongest independent components (ICs) that capture different modes of covariations in VBM and WM microstructure in addition to their relationships with age (A-B), PSQI (C-D) and WASO (E). Note that the x-axis shows the demeaned demographic (i.e., age) and behavioural (i.e., PSQI, WASO) values. The effect magnitude is represented by R^2^. The significant relationships between the IC subject loadings and demographic and behavioural measures are corrected for multiple comparisons set at p < 0.05.

Although the IC_1_ subject loadings supported that GM volumetric features contributed less to age-related changes in this component compared to WM microstructural features, both cortical (AG, SFG, STG, MFG, FP, lateral occipital cortex, parietal operculum cortex) and subcortical (caudate, thalamus) regions were involved in localised GM alterations (Figure 5). FA and MD maps revealed high degree of overlap in small clusters that correspond to the association (ILF, SLF), projection (ATR) and commissural (FMA, FMI) fibre bundles. Of note, the FA map showed ageing effects in other association (IFO and UF) fibre bundles whereas the MD map showed some unique changes in the limbic (CBH, FX) fibre bundles. On the other hand, the spatial map of IC_2_ showed widespread GM volumetric alterations with increasing age which were evident in the primary motor and somatosensory cortices, auditory cortex, limbic system, in addition to several prefrontal, language and visual related areas (Figure 5). Similarly, FA and MD shared considerable overlap in the tracts that form part of the limbic (CBH) and association (ILF, IFO, SLF) fibres. However, FA showed distinct age-related alterations in other limbic (FX), projection (FMA) and commissural (ATR, PTR) fibre bundles.

**Figure 5.**
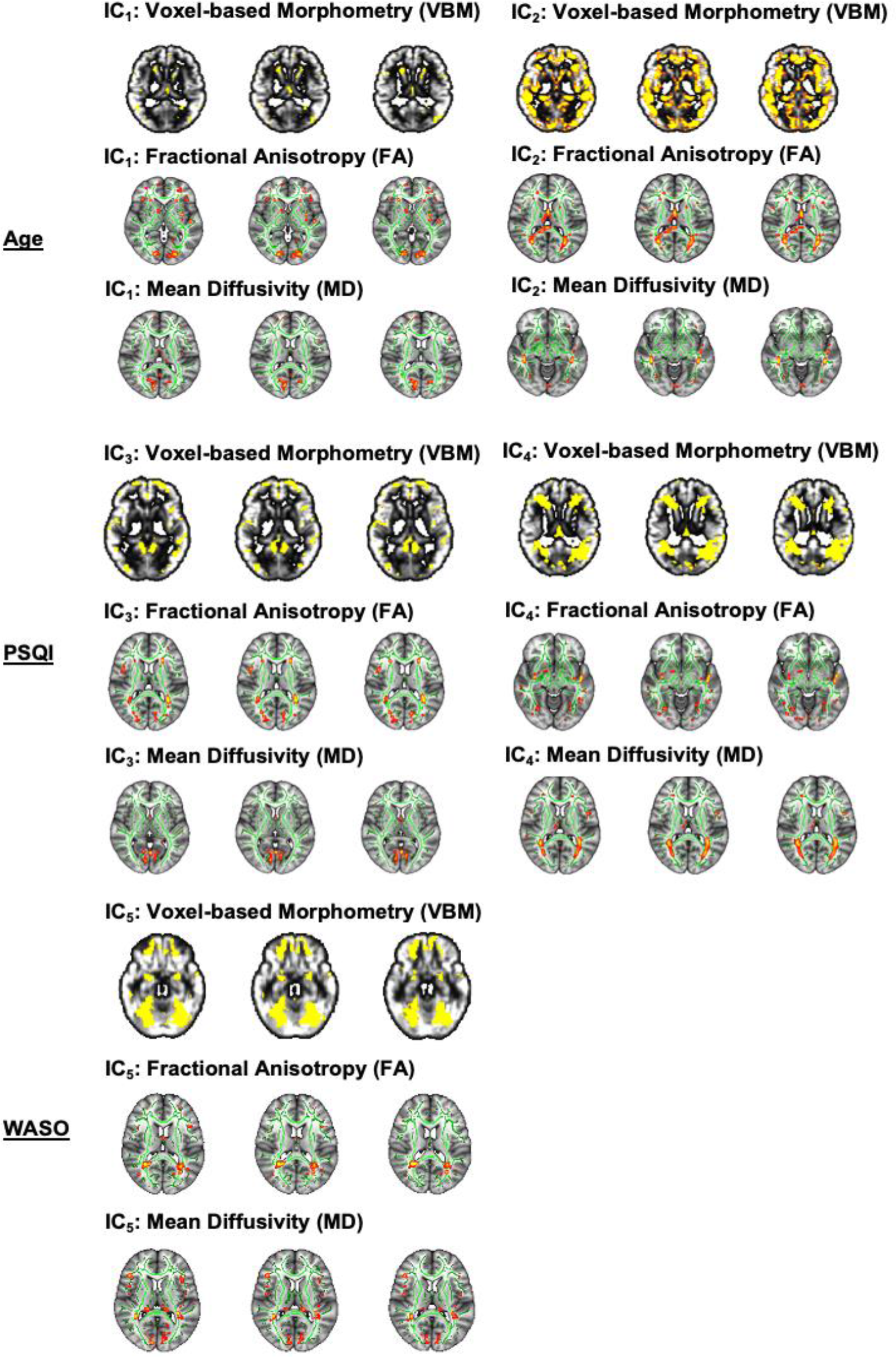
FLICA: Spatial maps of the independent components (ICs) that reveal GM volumetric and WM microstructural changes measured by FA and MD in Age (Top Panel), PSQI (Middle Panel) and WASO (Bottom Panel). The spatial maps are corrected for multiple comparisons set at p < 0.05.

### 3.6 FLICA: Covariation in GM and WM changes associated with sleep problems

Contrary to the unimodal TBSS and VBM analyses, multimodal FLICA analysis revealed associations between changes in GM structure and WM microstructure and sleep quality measures. FLICA revealed two multimodal components (i.e., IC_3_ and IC_4_) which captured interlinked GM and WM changes that were associated with the global sleep quality score, i.e., PSQI. IC_3_ showed a dominant mode of linear decrease in GM volumes and WM microstructure with poorer sleep quality (Figure 4C). PSQI explained 13% of the variance within IC_3_ (R^2^ = 0.13, p = 0.012 corrected). The linear relationship between the IC_3_ subject loadings and PSQI was mainly driven by GM volume (40%) followed by FA (31%) and MD (26%). The second PSQI-related component, IC_4_, shared two modes of linear and nonlinear covariation in regional GM and WM microstructure which were related to decreasing sleep quality (Figure 4D). The IC_4_ subject loadings showed a U-shape profile such that weaker loadings were associated with higher sleep quality (PSQI ≤ 5) whereas stronger loadings were related to a deterioration in sleep quality (PSQI > 5). While PSQI explained 10% of the linear variation within IC_4_ (R^2^ = 0.10, p = 0.024 corrected), it explained 24% of the U-shape variation within the same component (R^2^ = 0.24, p = 0.0059 corrected). Specifically, the linear and U-shape relationships between the IC_4_ subject loadings and PSQI were greatly dominated by morphological reductions (72%) followed by FA (15%) and MD (12%) alterations.

The spatial distribution of IC_3_ showed GM networks that have been previously associated with several cognitive domains such as language, sensorimotor, visual, emotion, attention and default mode functions (Figure 5). FA and MD maps demonstrated WM alterations with poor sleep quality in tracts that overlapped in the association (ILF, IFO) and projection (ATR) bundles. We also observed that FA captured distinct sleep-related effects in the FMA, AR, SLF and UF fibre bundles whereas MD showed some changes that were specific to the FX. Our data-driven approach also showed that the GM spatial map of IC_4_ overlapped with that of IC_3_ to a certain extent (Figure 5). Further, FA and MD revealed sleep-related alterations which overlapped in the association (ILF, IFO, SLF) and commissural (FMA) fibre bundles. Nonetheless, FA showed specific sleep-related changes in the UF and CBH fibre bundles whereas MD revealed non-overlapping alterations in the ATR.

Lastly, we observed a multimodal component (i.e., IC_5_) which revealed a significant nonlinear relationship with sleep fragmentation, i.e., WASO. IC_5_ exhibited a markedly inverted U-shape profile that reflects widespread GM and localised WM microstructural alterations with WASO (Figure 4E). The IC_5_ subject loadings displayed a nearly symmetrical inverted U-shape profile as a result of stronger loadings being associated with less sleep disturbance (WASO ≤ 36min) and weaker loadings relating to more frequent sleep disturbance (WASO > 36min). WASO explained 20% of the variance within IC_5_ (R^2^ = 0.20, p = 0.0017 corrected). The inverted U-shape relationship between the IC_5_ loading and WASO was predominantly driven by the GM volume (54%) followed by FA (27%) and MD (16%).

Spatially, the GM changes in IC_5_ were evident in the frontal, temporal and occipital areas including portions of the cerebellum (Figure 5). FA and MD maps showed significant WM alterations that overlapped in small clusters that form part of the association (ILF, IFO, SLF), projection (ATR) and commissural (FMA) fibre bundles. In addition, there were evidence of distinct sleep-related effects in the FA map that correspond to the limbic (CBH, FX) fibre bundles. Similarly, the MD map showed the presence of non-overlapping sleep-related effects in the PTR.

### 3.7 Brain Age Predictions

Using multimodal imaging features representing regional GM volumes and WM tract microstructure, we calculated the uncorrected (δ_b_), corrected linear (δ) and quadratic (δ_q_) estimates of brain age delta (Table 2). The magnitude of δ and δ_q_ showed that in the studied cohort, the estimated brain age (using the multimodal neuroimaging features) was on average approximately 2 years older than the chronological age. Both δ and 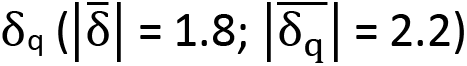 were notably smaller than 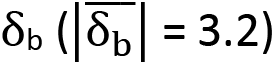, i.e., the biased/uncorrected estimate that is typically used in brain age prediction studies. Given that brain age delta corresponds to the residuals in the brain prediction model, the smaller δ and δ_q_ (which are orthogonal to age) improve the associations between the respective chronological age and predicted brain age (Y_L_ and Y_Q_ respectively) compared to the predicted brain age Y_B_ using δ_b_ (Table 2), as expected and shown before (Smith et al., 2019).

**Table 2.**
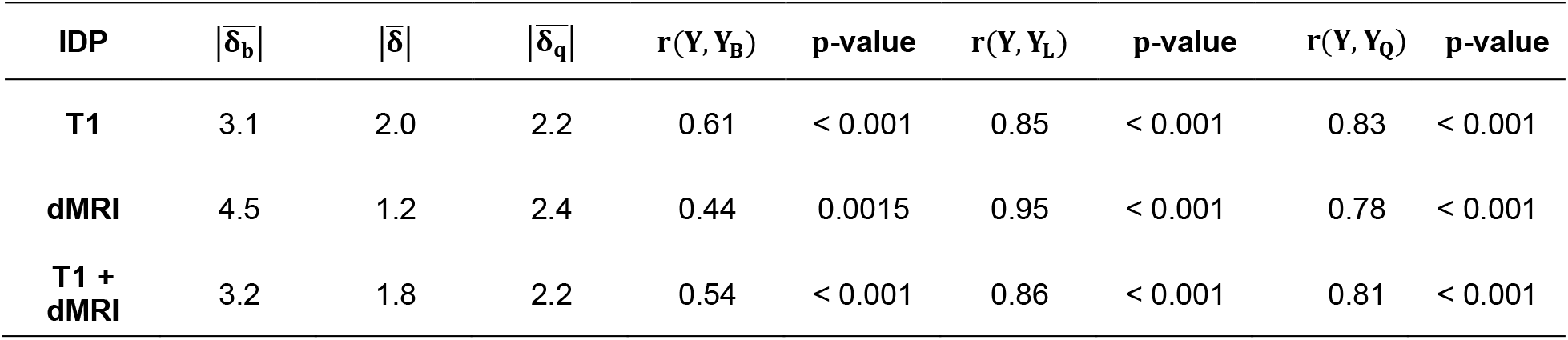
Brain Age Delta. Magnitude of the uncorrected (δ_b_), linearly-corrected (δ) and quadratically-corrected (δ_q_) estimates of delta obtained from unimodal (T1 or dMRI IDPs) and multimodal (T1+dMRI) IDPs, and the associations of chronological age (Y) with the uncorrected brain age prediction (Y_B_), linearly-corrected brain age prediction (Y_L_) and quadratically-corrected (Y_Q_) brain age prediction.

| IDP | $\vert \bar{\delta}_b\vert $ | $\vert \bar{\delta}\vert $ | $\vert \bar{\delta}_q\vert $ | $r(Y, Y_B)$ | p-value | $r(Y, Y_L)$ | p-value | $r(Y, Y_Q)$ | p-value |
| --- | --- | --- | --- | --- | --- | --- | --- | --- | --- |
| T1 | 3.1 | 2.0 | 2.2 | 0.61 | < 0.001 | 0.85 | < 0.001 | 0.83 | < 0.001 |
| dMRI | 4.5 | 1.2 | 2.4 | 0.44 | 0.0015 | 0.95 | < 0.001 | 0.78 | < 0.001 |
| T1 + dMRI | 3.2 | 1.8 | 2.2 | 0.54 | < 0.001 | 0.86 | < 0.001 | 0.81 | < 0.001 |

Brain age δ estimates using linear or quadratic corrections showed significant associations with poor sleep indices. Notably, there was no significant association between uncorrected δ_b_ with either PSQI or WASO.

When using unimodal (either T1 or dMRI) IDPs to estimate brain age delta, there was a significant relationship between the quadratically-corrected δ_q_ and PSQI (R = 0.30, p = 0.05 FDR) but only when the T1 IDPs were applied in the brain age prediction model (Figure 6A) (there was no significant relationship between δ_q_ and WASO (Figure 6B)).

**Figure 6.**
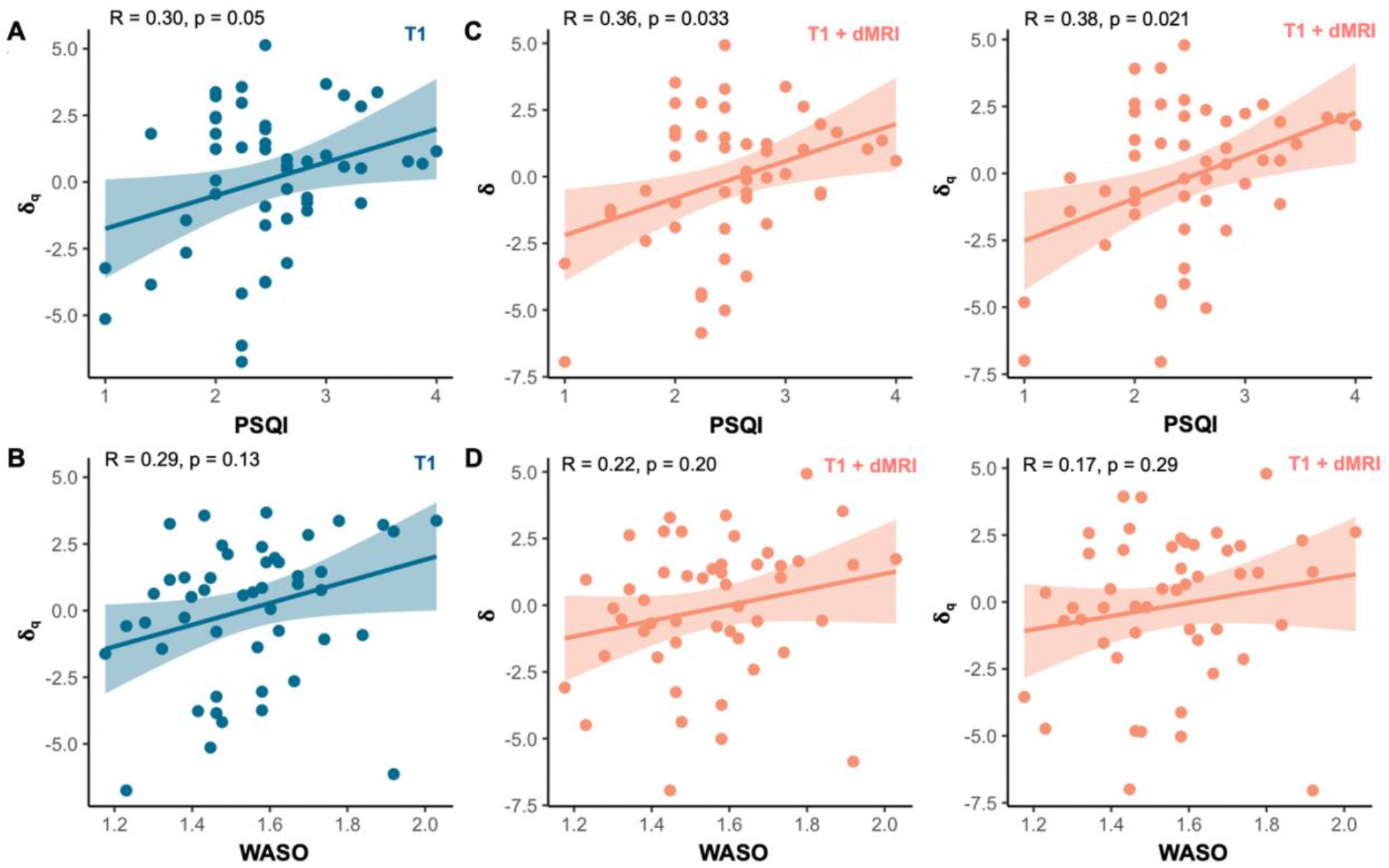
Relationships between brain age delta and sleep measures using the unimodal (T1) and multimodal (T1+dMRI) IDPs. **(A)** δ_q_ was significantly associated with PSQI using the unimodal T1 IDPs. **(B)**δ was not significantly associated with WASO after applying FDR correction when the unimodal T1 IDPs were used. **(C)** Both δ and δ_q_ were significantly related to PSQI with the multimodal (T1+dMRI) IDPs. **(D)** Neither δ nor δ_q_ was significantly associated with WASO when the multimodal (T1+dMRI) IDPs were used. All relationships between brain age delta and sleep measures are corrected for multiple comparisons set at p < 0.05. Note that the raw PSQI and WASO scores are square root and log-transformed, respectively (see section 2.2).

When using multimodal (T1+dMRI) IDPs to estimate brain age delta, the linearly-corrected δ estimate was significantly associated with PSQI (R = 0.36, p = 0.033 FDR) (Figure 6C). There was also a significant relationship between the quadratically-corrected δ_q_ and PSQI (R = 0.36, p = 0.021 FDR) when the multimodal IDPs were used (Figure 6C). Taken together, the results indicate that poor sleep quality is associated with accelerated brain ageing, i.e., brain age which is ~2 years older than its chronological age (neither δ nor δ_q_ was significantly associated with WASO (Figure 6D)).

## DISCUSSION

The ageing process, which results in structural brain deterioration and affects cognitive performance and daily functioning, is inevitable. However, not all older adults experience sharp cognitive decline. Some individuals undergo only gradual drop in cognitive functioning or even retain high levels of mental capacity throughout lifespan (Hayden et al., 2011). Substantial inter-individual differences in the pace of age-related brain changes underlying cognitive decline have been reported (for review see Cabeza et al., 2017; Eavani et al., 2018; Raz & Rodrigue, 2006). Consequently, disentangling the heterogeneity of brain ageing and factors underlying differential trajectories of normal versus pathological brain ageing is of high interest.

The current study explored volumetric and microstructural brain changes in healthy ageing and used estimates of brain age delta (difference between chronological and apparent brain age assessed using neuroimaging data) to investigate the associations of age-related brain changes with sleep quality and sleep fragmentation. Linked independent component analysis revealed a significant interlinked linear decrease of the global GM and WM microstructure with increasing age and sleep problems (both poor sleep quality and sleep fragmentation) as well as a degree of inter-individual variability in the observed age-related and poor sleep-related brain deterioration. These joint associations between brain structural and microstructural features with poor sleep indices were not evident using unimodal analyses.

Furthermore, brain age delta, estimated with linear and quadratic age-bias correction (Smith et al., 2019) from GM structural and WM microstructural neuroimaging features, revealed significant effects of poor sleep quality on the ageing brain. These associations were not evident when using the uncorrected (biased) estimates often used in brain-age prediction studies (for discussion see Le et al., 2018; Liang et al., 2019). Specifically, our findings demonstrated a two-year deviation above the chronological age (i.e., accelerated ageing) linked to sleep problems. We discuss our findings in context of prior research investigating the effects of sleep on age-related brain deterioration as well as research focusing on the use “brain age gap” as a biomarker of brain ageing.

The reported heterogeneity of age-related brain changes raises a possibility of maintaining more youthful brain later in life. While, both genetic and environmental influences likely account for variation in ageing, recent research strongly suggests that modifiable lifestyle factors, such as sleep, diet and physical activity, could hold the key to slowing down brain and cognitive ageing (for review see Wassenaar et al., 2019). As we get older waking up refreshed after a good night sleep becomes a challenge due to difficulties in maintaining or initiating sleep, fragmentation of sleep, increased daytime napping and changes in wake-sleep cycle (Miner & Kryger, 2017; Varma et al., 2019). The loss of good night sleep has a detrimental effect on overall health, reduces longevity and worsens cognitive performance (André et al., 2019; Gangwisch et al., 2008; Mattis & Sehgal, 2016). While older adults might have increased vulnerability to sleep problems and they often sleep less, the sleep recommendations for that age group are similar to these given to general adult population (7-9 hours per day; Hirshkowitz et al., 2015). Difficulties falling asleep, short sleep duration, excessive daytime sleepiness and napping can be regarded as modifiable behavioural sleep problems exacerbated by inadequate sleep hygiene, a set of behavioural practices, everyday habits and environmental factors required to achieve good night sleep (Lin et al., 2007). And sleep hygiene interventions have been shown to be effective in alleviating poor sleep in older adults, particularly increasing the efficiency of sleep and decreasing sleep fragmentation (e.g., Martin et al., 2017). Taken together, a growing body of evidence strongly suggests that reduced sleep time and poor sleep quality should no longer be viewed as intrinsically related to normal ageing but as potentially modifiable factors putting older adults at risk of cognitive decline and dementia (e.g., Ju et al., 2014; Keage et al., 2012; Lim et al., 2013; Wassenaar et al., 2019). While there is a substantial number of studies linking short sleep duration and/or poor sleep quality to grey matter atrophy and microstructural white matter changes, the evidence is inconsistent (Fjell et al., 2021; Lo et al., 2014; Ramos et al., 2014; Sexton et al., 2014; Sexton et al., 2020; Sexton et al., 2017; Spira et al., 2016; Yaffe et al., 2016; Zitser et al., 2020). Some of the reported discrepancies could be attributed to how sleep patterns are assessed, how brain changes are characterized and/or differences in studied population characteristics and/or sample size.

A recent large longitudinal study (613 participants), spanning over 28 years found no significant link between self-reported sleep duration and either grey matter or white matter microstructure (Zitser et al., 2020). However, it should be noted that Zitser and colleagues only collected longitudinal measures of sleep duration, while the brain changes were assessed at a single time-point. There are three potential explanations of these null results. Firstly, their study used unimodal analyses; we also could not find associations when performing unimodal analyses and only when probing the joint variance across modalities effects could be revealed. Analyses based on a single-item self-report of sleep duration in combination of single modality derived measures of GM or WM changes might not be sensitive enough to detect any brain deterioration beyond the effect of age itself. Secondly, it is plausible that not sleep duration per se but sleep quality or combination of these two affects brain changes in older adults without any diagnosis of sleep disorders. Thirdly, reliability of self-report of sleep duration as the discrepancies between objective and frequently overestimated self-reported measures of sleep duration, and their associations with health outcomes, are well documented (Lauderdale et al., 2008; McSorley et al., 2019; Silva et al., 2007).

The current study used a combination of objective (actigraphy based assessment of sleep fragmentation; WASO) and self-reported (PSQI questionnaire) measures of sleep to explore associations between sleep problems and age-related brain changes, hypothesizing that poor sleep accelerates brain ageing. We showed that self-reported but objectively measured sleep disruptions were associated with accelerated brain ageing. Specifically, global PSQI score indexing several common sleep dysfunctions (e.g., short sleep duration, sleep latency, poor sleep efficiency, sleep disturbances and daytime dysfunctions), was positively correlated with estimated brain age delta. Thus, indicating that “poorer” sleep is indeed linked to a larger “brain age gap” i.e., deviation from chronological age. Our findings suggest that subjective measures of sleep disruptions and/or composite measures of self-reported sleep problems are more sensitive to detect the effect of sleep on the ageing brain as compared to single item self-reported measures. Alternatively, it is plausible that overall poor sleep quality rather than sleep duration itself (see Zitser et al., 2020) has a substantially greater effect on brain deterioration in neurotypical older adults. As the observed heterogeneity of brain ageing can be attributed to both the large variability in the pace of ageing and to a wide range of onset ages for ageing process, the detection of any brain deterioration beyond the effect of age itself, e.g., the impact of sleep, might be challenging. This perhaps explains the discrepancies between our findings and some of the prior studies examining the effect of sleep on the ageing brain, which all independently examined patterns of either gray or white matter changes estimated from a single MRI modality. Furthermore, to increase sensitivity, we used here measures derived from both structural and diffusion MRI to examine interlinked GM and WM changes as well as estimated deviation from chronological age (brain age delta; Smith et al., 2019) to investigate the link between age-related brain changes and poor sleep in a group of neurotypical elderly participants. We explored that further using exploratory FLICA method (Douaud et al., 2014; Groves et al., 2011).

The FLICA approach applied here demonstrated an interlinked global GM and WM decrease with increasing age and a variability in the observed age-related brain deterioration in the studied group of elderly participants. Importantly, FLICA revealed interlinked GM and WM changes driven by poor sleep as assessed by PSQI. This is in line with both single modality and multimodal estimates of brain age delta, which proved to be a sensitive biomarker of accelerated brain ageing linked to poor sleep quality. FLICA also identified a multimodal component associated with sleep fragmentation (WASO). The U-shape versus linear profile of the unique multimodal components revealed by FLICA are result of differential contribution of GM volume, FA and MD to each individual component (Douaud et al., 2014; Groves et al., 2011). It should be noted here that in contrast to FLICA, commonly used unimodal approaches such as VBM or TBSS, separately using measures of either GM or WM changes, failed to detect any links between poor sleep and brain deterioration in the examined group of elderly participants. Thus, these results strongly indicate that multimodal analysis increases sensitivity when assessing association between sleep problems and brain ageing.

“Brain age gap” estimates are increasingly being used as a biomarker of brain’s health signaling increased risk of brain deterioration and predictors of progression from mild cognitive impairment to dementia (Cole & Franke, 2017; Gaser et al., 2013). This method has been also applied to predict cognitive functioning in non-demented older adults and to identify lifestyle factors associated with maintaining a more youthful brain in old age (Boyle et al., 2021; Steffener et al., 2016). While many, especially earlier, “brain age gap” studies used estimates based on a single MRI modality (for review see Cole & Franke, 2017; Franke & Gasser, 2019), more recent approaches employed multimodal datasets (Cole, 2020; Smith et al., 2020; Smith et al., 2019). Here, we used a recently developed multimodal technique to calculate unbiased estimates of brain age delta (see Smith et al., 2019), in order to explore the effects of sleep on the ageing brain, stipulating that poor sleep would accelerate brain ageing. As proposed by Smith and colleagues, this approach enables, to remove typical biases affecting brain-age estimates (and lead to overestimation in younger and underestimation in older individuals; for discussion see Le et al., 2018; Liang et al., 2019); see also Beheshti et al., 2019) for similar approach) and increases sensitivity of associations with lifestyle/socio-demographic variables. Indeed, we found that the corrected (i.e., removal of age-related bias) brain age delta estimates outperformed the uncorrected (biased) estimates in their predictive power and revealed associations with poor sleep indices (PSQI), not evident when the uncorrected delta estimates were used. In the studied group of neurotypical older adults we found two years deviation above the chronological age, which was correlated with poor sleep. A two-year deviation from chronological age might seem relatively small. However, to put our findings in a broader perspective, it should be noted that a recent large epidemiological study identified a four-year brain age gap to be associated with dementia and predictive of low cognitive functioning (Kaufmann et al., 2019).

Taking into account a recent evidence that a few years deviation from normative brain ageing is one of the hallmarks of dementia (Kaufmann et al., 2019), we conclude that sleep problems in healthy older adults should be considered a modifiable risk factor for dementia. Thus, our findings also point to the aptitude of behavioural intervention to combat the effects of poor sleep on the ageing brain. However, it should be noted that any conclusions drawn from our findings are limited by cross-sectional design and thus further longitudinal studies, preferably based on multimodal approaches are needed.

## ACKNOWLEDGMENTS

We warmly thank the volunteers from the School of Psychology panel and the Birmingham 1000 Elders group for participation in this study. This work was supported by the Birmingham-Nottingham Strategic Collaboration Fund (BNSCF336 to SNS and MC) and by a Wellcome Trust Institutional Strategic Support Fund critical data award (204846/Z/16/Z to MC). SS was also supported by the Wellcome Trust (217266/Z/19/Z) and MC was also supported by a BRIDGE (Birmingham-Illinois Partnership for Discovery, Engagement and Education) Fellowship.

## DATA AVAILABILITY STATEMENT

Participantdata will be made available to researchers upon request.

## ETHICS

The study was approved by the University of Birmingham Ethical Review Committee. All participants provided written informed consent in agreement with approved ethics protocol (ERN_110429AP66).

## CONFLICT OF INTEREST

The authors declare no potential conflict of interest.

## Notes

### Competing Interest Statement

The authors have declared no competing interest.

## REFERENCES

Alfaro-Almagro, F., Jenkinson, M., Bangerter, N. K., Andersson, J. L. R., Griffanti, L., Douaud, G., Sotiropoulos, S. N., Jbabdi, S., Hernandez-Fernandez, M., Vallee, E., Vidaurre, D., Webster, M., McCarthy, P., Rorden, C., Daducci, A., Alexander, D. C., Zhang, H., Dragonu, I., Matthews, P. M., Miller, K. L., & Smith, S. M. (2018). Image processing and Quality Control for the first 10,000 brain imaging datasets from UK Biobank. Neuroimage, 166, 400–424. https://doi.org/10.1016/j.neuroimage.2017.10.034

Andersson, J. L., Skare, S., & Ashburner, J. (2003). How to correct susceptibility distortions in spin-echo echo-planar images: application to diffusion tensor imaging. Neuroimage, 20(2), 870–888. https://doi.org/10.1016/s1053-8119(03)00336-7

Andersson, J. L. R., Jenkinson, M., & Smith, S. (2019). High resolution nonlinear registration with simultaneous modelling of intensities. bioRxiv, 646802. https://doi.org/10.1101/646802

Andersson, J. L. R., & Sotiropoulos, S. N. (2016). An integrated approach to correction for off-resonance effects and subject movement in diffusion MR imaging. Neuroimage, 125, 1063–1078. https://doi.org/10.1016/j.neuroimage.2015.10.019

André, C., Tomadesso, C., de Flores, R., Branger, P., Rehel, S., Mézenge, F., Landeau, B., Sayette, V. d. l., Eustache, F., Chételat, G., & Rauchs, G. (2019). Brain and cognitive correlates of sleep fragmentation in elderly subjects with and without cognitive deficits. Alzheimer’s & dementia (Amsterdam, Netherlands), 11, 142–150. https://doi.org/10.1016/j.dadm.2018.12.009

Ashburner, J., & Friston, K. J. (2000). Voxel-based morphometry--the methods. Neuroimage, 11(6 Pt 1), 805–821. https://doi.org/10.1006/nimg.2000.0582

Barrick, T. R., Charlton, R. A., Clark, C. A., & Markus, H. S. (2010). White matter structural decline in normal ageing: a prospective longitudinal study using tract-based spatial statistics. Neuroimage, 51(2), 565–577. https://doi.org/10.1016/j.neuroimage.2010.02.033

Basser, P. J., Mattiello, J., & Lebihan, D. (1994). Estimation of the Effective Self-Diffusion Tensor from the NMR Spin Echo. Journal of Magnetic Resonance, Series B, 103(3), 247–254. https://doi.org/10.1006/jmrb.1994.1037

Beheshti, I., Nugent, S., Potvin, O., & Duchesne, S. (2019). Bias-adjustment in neuroimaging-based brain age frameworks: A robust scheme. NeuroImage. Clinical, 24, 102063–102063. https://doi.org/10.1016/j.nicl.2019.102063

Behrens, T. E., Berg, H. J., Jbabdi, S., Rushworth, M. F., & Woolrich, M. W. (2007). Probabilistic diffusion tractography with multiple fibre orientations: What can we gain? Neuroimage, 34(1), 144–155. https://doi.org/10.1016/j.neuroimage.2006.09.018

Boyle, R., Jollans, L., Rueda-Delgado, L. M., Rizzo, R., Yener, G. G., McMorrow, J. P., Knight, S. P., Carey, D., Robertson, I. H., Emek-Savaş, D. D., Stern, Y., Kenny, R. A., & Whelan, R. (2021). Brain-predicted age difference score is related to specific cognitive functions: a multi-site replication analysis. Brain Imaging and Behavior, 15(1), 327–345. https://doi.org/10.1007/s11682-020-00260-3

Burzynska, A. Z., Preuschhof, C., Bäckman, L., Nyberg, L., Li, S. C., Lindenberger, U., & Heekeren, H. R. (2010). Age-related differences in white matter microstructure: region-specific patterns of diffusivity. Neuroimage, 49(3), 2104–2112. https://doi.org/10.1016/j.neuroimage.2009.09.041

Buysse, D. J., Reynolds, C. F., 3rd, Monk, T. H., Berman, S. R., & Kupfer, D. J. (1989). The Pittsburgh Sleep Quality Index: a new instrument for psychiatric practice and research. Psychiatry Res, 28(2), 193–213. https://doi.org/10.1016/0165-1781(89)90047-4

Buysse, D. J., Reynolds, C. F., 3rd, Monk, T. H., Hoch, C. C., Yeager, A. L., & Kupfer, D. J. (1991). Quantification of subjective sleep quality in healthy elderly men and women using the Pittsburgh Sleep Quality Index (PSQI). Sleep, 14(4), 331–338.

Cabeza, R., Albert, M., Belleville, S., Craik, F. I. M., Duarte, A., Grady, C. L., Lindenberger, U., Nyberg, L., Park, D. C., Reuter-Lorenz, P. A., Rugg, M. D., Steffener, J., & Rajah, M. N. (2018). Maintenance, reserve and compensation: the cognitive neuroscience of healthy ageing. Nat Rev Neurosci, 19(11), 701–710. https://doi.org/10.1038/s41583-018-0068-2

Cabeza, R., Nyberg, L., & Park, D. C. (2017). Cognitive neuroscience of aging: linking cognitive and cerebral aging (Second edition. ed.) [still image]. Oxford University Press.

Carpenter, J. S., & Andrykowski, M. A. (1998). Psychometric evaluation of the Pittsburgh Sleep Quality Index. J Psychosom Res, 45(1), 5–13. https://doi.org/10.1016/s0022-3999(97)00298-5

Cole, J. H. (2020). Multimodality neuroimaging brain-age in UK biobank: relationship to biomedical, lifestyle, and cognitive factors. Neurobiol Aging, 92, 34–42. https://doi.org/10.1016/j.neurobiolaging.2020.03.014

Cole, J. H., & Franke, K. (2017). Predicting Age Using Neuroimaging: Innovative Brain Ageing Biomarkers. Trends Neurosci, 40(12), 681–690. https://doi.org/10.1016/j.tins.2017.10.001

Cole, J. H., Poudel, R. P. K., Tsagkrasoulis, D., Caan, M. W. A., Steves, C., Spector, T. D., & Montana, G. (2017). Predicting brain age with deep learning from raw imaging data results in a reliable and heritable biomarker. Neuroimage, 163, 115–124. https://doi.org/10.1016/j.neuroimage.2017.07.059

Dale, A. M., Fischl, B., & Sereno, M. I. (1999). Cortical surface-based analysis. I. Segmentation and surface reconstruction. Neuroimage, 9(2), 179–194. https://doi.org/10.1006/nimg.1998.0395

Douaud, G., Groves, A. R., Tamnes, C. K., Westlye, L. T., Duff, E. P., Engvig, A., Walhovd, K. B., James, A., Gass, A., Monsch, A. U., Matthews, P. M., Fjell, A. M., Smith, S. M., & Johansen-Berg, H. (2014). A common brain network links development, aging, and vulnerability to disease. Proceedings of the National Academy of Sciences, 111(49), 17648–17653. https://doi.org/10.1073/pnas.1410378111

Douaud, G., Smith, S., Jenkinson, M., Behrens, T., Johansen-Berg, H., Vickers, J., James, S., Voets, N., Watkins, K., Matthews, P. M., & James, A. (2007). Anatomically related grey and white matter abnormalities in adolescent-onset schizophrenia. Brain, 130(9), 2375–2386. https://doi.org/10.1093/brain/awm184

Eavani, H., Habes, M., Satterthwaite, T. D., An, Y., Hsieh, M. K., Honnorat, N., Erus, G., Doshi, J., Ferrucci, L., Beason-Held, L. L., Resnick, S. M., & Davatzikos, C. (2018). Heterogeneity of structural and functional imaging patterns of advanced brain aging revealed via machine learning methods. Neurobiol Aging, 71, 41–50. https://doi.org/10.1016/j.neurobiolaging.2018.06.013

Fabiani, M. (2012). It was the best of times, it was the worst of times: a psychophysiologist’s view of cognitive aging. Psychophysiology, 49(3), 283–304. https://doi.org/10.1111/j.1469-8986.2011.01331.x

Fischl, B. (2012). FreeSurfer. Neuroimage, 62(2), 774–781. https://doi.org/10.1016/j.neuroimage.2012.01.021

Fischl, B., & Dale, A. M. (2000). Measuring the thickness of the human cerebral cortex from magnetic resonance images. Proc Natl Acad Sci U S A, 97(20), 11050–11055. https://doi.org/10.1073/pnas.200033797

Fjell, A. M., Sørensen, Ø., Amlien, I. K., Bartrés-Faz, D., Brandmaier, A. M., Buchmann, N., Demuth, I., Drevon, C. A., Düzel, S., Ebmeier, K. P., Ghisletta, P., Idland, A.-V., Kietzmann, T. C., Kievit, R. A., Kühn, S., Lindenberger, U., Magnussen, F., Macià, D., Mowinckel, A. M., Nyberg, L., Sexton, C. E., Solé-Padullés, C., Pudas, S., Roe, J. M., Sederevicius, D., Suri, S., Vidal-Piñeiro, D., Wagner, G., Watne, L. O., Westerhausen, R., Zsoldos, E., & Walhovd, K. B. (2021). Poor Self-Reported Sleep is Related to Regional Cortical Thinning in Aging but not Memory Decline—Results From the Lifebrain Consortium. Cerebral Cortex, 31(4), 1953–1969. https://doi.org/10.1093/cercor/bhaa332

Franke, K., & Gaser, C. (2019). Ten Years of BrainAGE as a Neuroimaging Biomarker of Brain Aging: What Insights Have We Gained? Front Neurol, 10, 789. https://doi.org/10.3389/fneur.2019.00789

Gangwisch, J. E., Heymsfield, S. B., Boden-Albala, B., Buijs, R. M., Kreier, F., Opler, M. G., Pickering, T. G., Rundle, A. G., Zammit, G. K., & Malaspina, D. (2008). Sleep duration associated with mortality in elderly, but not middle-aged, adults in a large US sample. Sleep, 31(8), 1087–1096.

Gaser, C., Franke, K., Klöppel, S., Koutsouleris, N., Sauer, H., & Alzheimer’s Disease Neuroimaging, I. (2013). BrainAGE in Mild Cognitive Impaired Patients: Predicting the Conversion to Alzheimer’s Disease. PLOS ONE, 8(6), e67346. https://doi.org/10.1371/journal.pone.0067346

Gentili, A., Weiner, D. K., Kuchibhatla, M., & Edinger, J. D. (1995). Test-retest reliability of the Pittsburgh sleep quality index in nursing home residents. J Am Geriatr Soc, 43(11), 1317–1318. https://doi.org/10.1111/j.1532-5415.1995.tb07415.x

Giorgio, A., Santelli, L., Tomassini, V., Bosnell, R., Smith, S., De Stefano, N., & Johansen-Berg, H. (2010). Age-related changes in grey and white matter structure throughout adulthood. Neuroimage, 51(3), 943–951. https://doi.org/10.1016/j.neuroimage.2010.03.004

Good, C. D., Johnsrude, I. S., Ashburner, J., Henson, R. N., Friston, K. J., & Frackowiak, R. S. (2001). A voxel-based morphometric study of ageing in 465 normal adult human brains. Neuroimage, 14(1 Pt 1), 21–36. https://doi.org/10.1006/nimg.2001.0786

Groves, A. R., Beckmann, C. F., Smith, S. M., & Woolrich, M. W. (2011). Linked independent component analysis for multimodal data fusion. Neuroimage, 54(3), 2198–2217. https://doi.org/10.1016/j.neuroimage.2010.09.073

Groves, A. R., Smith, S. M., Fjell, A. M., Tamnes, C. K., Walhovd, K. B., Douaud, G., Woolrich, M. W., & Westlye, L. T. (2012). Benefits of multi-modal fusion analysis on a large-scale dataset: Life-span patterns of inter-subject variability in cortical morphometry and white matter microstructure. Neuroimage, 63(1), 365–380. https://doi.org/10.1016/j.neuroimage.2012.06.038

Hayden, K. M., Reed, B. R., Manly, J. J., Tommet, D., Pietrzak, R. H., Chelune, G. J., Yang, F. M., Revell, A. J., Bennett, D. A., & Jones, R. N. (2011). Cognitive decline in the elderly: an analysis of population heterogeneity. Age Ageing, 40(6), 684–689. https://doi.org/10.1093/ageing/afr101

Hirshkowitz, M., Whiton, K., Albert, S. M., Alessi, C., Bruni, O., DonCarlos, L., Hazen, N., Herman, J., Katz, E. S., Kheirandish-Gozal, L., Neubauer, D. N., O’Donnell, A. E., Ohayon, M., Peever, J., Rawding, R., Sachdeva, R. C., Setters, B., Vitiello, M. V., Ware, J. C., & Adams Hillard, P. J. (2015). National Sleep Foundation’s sleep time duration recommendations: methodology and results summary. Sleep Health, 1(1), 40–43. https://doi.org/10.1016/j.sleh.2014.12.010

Ju, Y.-E. S., Lucey, B. P., & Holtzman, D. M. (2014). Sleep and Alzheimer disease pathology--a bidirectional relationship. Nature reviews. Neurology, 10(2), 115–119. https://doi.org/10.1038/nrneurol.2013.269

Kaufmann, T., van der Meer, D., Doan, N. T., Schwarz, E., Lund, M. J., Agartz, I., Alnæs, D., Barch, D. M., Baur-Streubel, R., Bertolino, A., Bettella, F., Beyer, M. K., Bøen, E., Borgwardt, S., Brandt, C. L., Buitelaar, J., Celius, E. G., Cervenka, S., Conzelmann, A., Córdova-Palomera, A., Dale, A. M., de Quervain, D. J. F., Di Carlo, P., Djurovic, S., Dørum, E. S., Eisenacher, S., Elvsåshagen, T., Espeseth, T., Fatouros-Bergman, H., Flyckt, L., Franke, B., Frei, O., Haatveit, B., Håberg, A. K., Harbo, H. F., Hartman, C. A., Heslenfeld, D., Hoekstra, P. J., Høgestøl, E. A., Jernigan, T. L., Jonassen, R., Jönsson, E. G., Farde, L., Flyckt, L., Engberg, G., Erhardt, S., Fatouros-Bergman, H., Cervenka, S., Schwieler, L., Piehl, F., Agartz, I., Collste, K., Victorsson, P., Malmqvist, A., Hedberg, M., Orhan, F., Kirsch, P., Kłoszewska, I., Kolskår, K. K., Landrø, N. I., Le Hellard, S., Lesch, K.-P., Lovestone, S., Lundervold, A., Lundervold, A. J., Maglanoc, L. A., Malt, U. F., Mecocci, P., Melle, I., Meyer-Lindenberg, A., Moberget, T., Norbom, L. B., Nordvik, J. E., Nyberg, L., Oosterlaan, J., Papalino, M., Papassotiropoulos, A., Pauli, P., Pergola, G., Persson, K., Richard, G., Rokicki, J., Sanders, A.-M., Selbæk, G., Shadrin, A. A., Smeland, O. B., Soininen, H., Sowa, P., Steen, V. M., Tsolaki, M., Ulrichsen, K. M., Vellas, B., Wang, L., Westman, E., Ziegler, G. C., Zink, M., Andreassen, O. A., Westlye, L. T., & Karolinska Schizophrenia, P. (2019). Common brain disorders are associated with heritable patterns of apparent aging of the brain. Nature Neuroscience, 22(10), 1617–1623. https://doi.org/10.1038/s41593-019-0471-7

Keage, H. A., Banks, S., Yang, K. L., Morgan, K., Brayne, C., & Matthews, F. E. (2012). What sleep characteristics predict cognitive decline in the elderly? Sleep Med, 13(7), 886–892. https://doi.org/10.1016/j.sleep.2012.02.003

Lauderdale, D. S., Knutson, K. L., Yan, L. L., Liu, K., & Rathouz, P. J. (2008). Self-reported and measured sleep duration: how similar are they? Epidemiology, 19(6), 838–845. https://doi.org/10.1097/EDE.0b013e318187a7b0

Le, T. T., Kuplicki, R. T., McKinney, B. A., Yeh, H.-W., Thompson, W. K., Paulus, M. P., , T. I., Aupperle, R. L., Bodurka, J., Cha, Y.-H., Feinstein, J. S., Khalsa, S. S., Savitz, J., Simmons, W. K., & Victor, T. A. (2018). A Nonlinear Simulation Framework Supports Adjusting for Age When Analyzing BrainAGE [Methods]. Frontiers in Aging Neuroscience, 10(317). https://doi.org/10.3389/fnagi.2018.00317

Lemaitre, H., Goldman, A. L., Sambataro, F., Verchinski, B. A., Meyer-Lindenberg, A., Weinberger, D. R., & Mattay, V. S. (2012). Normal age-related brain morphometric changes: nonuniformity across cortical thickness, surface area and gray matter volume? Neurobiol Aging, 33(3), 617.e611–619. https://doi.org/10.1016/j.neurobiolaging.2010.07.013

Liang, H., Zhang, F., & Niu, X. (2019). Investigating systematic bias in brain age estimation with application to post-traumatic stress disorders. Hum Brain Mapp, 40(11), 3143–3152. https://doi.org/10.1002/hbm.24588

Liem, F., Varoquaux, G., Kynast, J., Beyer, F., Kharabian Masouleh, S., Huntenburg, J. M., Lampe, L., Rahim, M., Abraham, A., Craddock, R. C., Riedel-Heller, S., Luck, T., Loeffler, M., Schroeter, M. L., Witte, A. V., Villringer, A., & Margulies, D. S. (2017). Predicting brain-age from multimodal imaging data captures cognitive impairment. Neuroimage, 148, 179–188. https://doi.org/10.1016/j.neuroimage.2016.11.005

Lim, A. S., Kowgier, M., Yu, L., Buchman, A. S., & Bennett, D. A. (2013). Sleep Fragmentation and the Risk of Incident Alzheimer’s Disease and Cognitive Decline in Older Persons. Sleep, 36(7), 1027–1032. https://doi.org/10.5665/sleep.2802

Lin, S. C., Cheng, C., Yang, C.-M., Hsui, S. C., & ©åò. (2007). Psychometric properties of the sleep hygiene practice scale. Sleep, 30, A262.

Liu, J., Pearlson, G., Windemuth, A., Ruano, G., Perrone-Bizzozero, N. I., & Calhoun, V. (2009). Combining fMRI and SNP data to investigate connections between brain function and genetics using parallel ICA. Hum Brain Mapp, 30(1), 241–255. https://doi.org/10.1002/hbm.20508

Lo, J. C., Loh, K. K., Zheng, H., Sim, S. K., & Chee, M. W. (2014). Sleep duration and age-related changes in brain structure and cognitive performance. Sleep, 37(7), 1171–1178. https://doi.org/10.5665/sleep.3832

Martin, J. L., Song, Y., Hughes, J., Jouldjian, S., Dzierzewski, J. M., Fung, C. H., Rodriguez Tapia, J. C., Mitchell, M. N., & Alessi, C. A. (2017). A Four-Session Sleep Intervention Program Improves Sleep for Older Adult Day Health Care Participants: Results of a Randomized Controlled Trial. Sleep, 40(8). https://doi.org/10.1093/sleep/zsx079

Mattis, J., & Sehgal, A. (2016). Circadian Rhythms, Sleep, and Disorders of Aging. Trends Endocrinol Metab, 27(4), 192–203. https://doi.org/10.1016/j.tem.2016.02.003

McGinnis, S. M., Brickhouse, M., Pascual, B., & Dickerson, B. C. (2011). Age-related changes in the thickness of cortical zones in humans. Brain Topogr, 24(3-4), 279–291. https://doi.org/10.1007/s10548-011-0198-6

McSorley, V. E., Bin, Y. S., & Lauderdale, D. S. (2019). Associations of Sleep Characteristics With Cognitive Function and Decline Among Older Adults. Am J Epidemiol, 188(6), 1066–1075. https://doi.org/10.1093/aje/kwz037

Miller, K. L., Alfaro-Almagro, F., Bangerter, N. K., Thomas, D. L., Yacoub, E., Xu, J., Bartsch, A. J., Jbabdi, S., Sotiropoulos, S. N., Andersson, J. L. R., Griffanti, L., Douaud, G., Okell, T. W., Weale, P., Dragonu, I., Garratt, S., Hudson, S., Collins, R., Jenkinson, M., Matthews, P. M., & Smith, S. M. (2016). Multimodal population brain imaging in the UK Biobank prospective epidemiological study. Nature Neuroscience, 19(11), 1523–1536. https://doi.org/10.1038/nn.4393

Miner, B., & Kryger, M. H. (2017). Sleep in the Aging Population. Sleep Med Clin, 12(1), 31–38. https://doi.org/10.1016/j.jsmc.2016.10.008

Patenaude, B., Smith, S. M., Kennedy, D. N., & Jenkinson, M. (2011). A Bayesian model of shape and appearance for subcortical brain segmentation. Neuroimage, 56(3), 907–922. https://doi.org/10.1016/j.neuroimage.2011.02.046

Ramos, A. R., Dong, C., Rundek, T., Elkind, M. S., Boden-Albala, B., Sacco, R. L., & Wright, C. B. (2014). Sleep duration is associated with white matter hyperintensity volume in older adults: the Northern Manhattan Study. J Sleep Res, 23(5), 524–530. https://doi.org/10.1111/jsr.12177

Raz, N., & Rodrigue, K. M. (2006). Differential aging of the brain: patterns, cognitive correlates and modifiers. Neurosci Biobehav Rev, 30(6), 730–748. https://doi.org/10.1016/j.neubiorev.2006.07.001

Sexton, C. E., Storsve, A. B., Walhovd, K. B., Johansen-Berg, H., & Fjell, A. M. (2014). Poor sleep quality is associated with increased cortical atrophy in community-dwelling adults. Neurology, 83(11), 967–973. https://doi.org/10.1212/wnl.0000000000000774

Sexton, C. E., Sykara, K., Karageorgiou, E., Zitser, J., Rosa, T., Yaffe, K., & Leng, Y. (2020). Connections Between Insomnia and Cognitive Aging. Neuroscience Bulletin, 36(1), 77–84. https://doi.org/10.1007/s12264-019-00401-9

Sexton, C. E., Zsoldos, E., Filippini, N., Griffanti, L., Winkler, A., Mahmood, A., Allan, C. L., Topiwala, A., Kyle, S. D., Spiegelhalder, K., Singh-Manoux, A., Kivimaki, M., Mackay, C. E., Johansen-Berg, H., & Ebmeier, K. P. (2017). Associations between self-reported sleep quality and white matter in community-dwelling older adults: A prospective cohort study. Hum Brain Mapp, 38(11), 5465–5473. https://doi.org/10.1002/hbm.23739

Silva, G. E., Goodwin, J. L., Sherrill, D. L., Arnold, J. L., Bootzin, R. R., Smith, T., Walsleben, J. A., Baldwin, C. M., & Quan, S. F. (2007). Relationship between reported and measured sleep times: the sleep heart health study (SHHS). Journal of clinical sleep medicine: JCSM: official publication of the American Academy of Sleep Medicine, 3(6), 622–630. https://pubmed.ncbi.nlm.nih.gov/17993045 https://www.ncbi.nlm.nih.gov/pmc/articles/PMC2045712/

Smith, S. M., Elliott, L. T., Alfaro-Almagro, F., McCarthy, P., Nichols, T. E., Douaud, G., & Miller, K. L. (2020). Brain aging comprises many modes of structural and functional change with distinct genetic and biophysical associations. eLife, 9, e52677. https://doi.org/10.7554/eLife.52677

Smith, S. M., Jenkinson, M., Johansen-Berg, H., Rueckert, D., Nichols, T. E., Mackay, C. E., Watkins, K. E., Ciccarelli, O., Cader, M. Z., Matthews, P. M., & Behrens, T. E. (2006). Tract-based spatial statistics: voxelwise analysis of multi-subject diffusion data. Neuroimage, 31(4), 1487–1505. https://doi.org/10.1016/j.neuroimage.2006.02.024

Smith, S. M., & Nichols, T. E. (2009). Threshold-free cluster enhancement: addressing problems of smoothing, threshold dependence and localisation in cluster inference. Neuroimage, 44(1), 83–98. https://doi.org/10.1016/j.neuroimage.2008.03.061

Smith, S. M., Vidaurre, D., Alfaro-Almagro, F., Nichols, T. E., & Miller, K. L. (2019). Estimation of brain age delta from brain imaging. Neuroimage, 200, 528–539. https://doi.org/10.1016/j.neuroimage.2019.06.017

Spira, A. P., Gonzalez, C. E., Venkatraman, V. K., Wu, M. N., Pacheco, J., Simonsick, E. M., Ferrucci, L., & Resnick, S. M. (2016). Sleep Duration and Subsequent Cortical Thinning in Cognitively Normal Older Adults. Sleep, 39(5), 1121–1128. https://doi.org/10.5665/sleep.5768

Steffener, J., Habeck, C., O’Shea, D., Razlighi, Q., Bherer, L., & Stern, Y. (2016). Differences between chronological and brain age are related to education and self-reported physical activity. Neurobiol Aging, 40, 138–144. https://doi.org/10.1016/j.neurobiolaging.2016.01.014

Stern, Y., Arenaza-Urquijo, E. M., Bartrés-Faz, D., Belleville, S., Cantilon, M., Chetelat, G., Ewers, M., Franzmeier, N., Kempermann, G., Kremen, W. S., Okonkwo, O., Scarmeas, N., Soldan, A., Udeh-Momoh, C., Valenzuela, M., Vemuri, P., & Vuoksimaa, E. (2020). Whitepaper: Defining and investigating cognitive reserve, brain reserve, and brain maintenance. Alzheimers Dement, 16(9), 1305–1311. https://doi.org/10.1016/j.jalz.2018.07.219

Varma, P., Jackson, M. L., & Meaklim, H. (2019). Dreaming of the good old days: sleep in older adults. Journal of Pharmacy Practice and Research, 49(3), 209–211. https://doi.org/10.1002/jppr.1578

Warrington, S., Bryant, K. L., Khrapitchev, A. A., Sallet, J., Charquero-Ballester, M., Douaud, G., Jbabdi, S., Mars, R. B., & Sotiropoulos, S. N. (2020). XTRACT - Standardised protocols for automated tractography in the human and macaque brain. Neuroimage, 217, 116923. https://doi.org/10.1016/j.neuroimage.2020.116923

Wassenaar, T. M., Yaffe, K., van der Werf, Y. D., & Sexton, C. E. (2019). Associations between modifiable risk factors and white matter of the aging brain: insights from diffusion tensor imaging studies. Neurobiol Aging, 80, 56–70. https://doi.org/10.1016/j.neurobiolaging.2019.04.006

Westlye, L. T., Walhovd, K. B., Dale, A. M., Bjørnerud, A., Due-Tønnessen, P., Engvig, A., Grydeland, H., Tamnes, C. K., Ostby, Y., & Fjell, A. M. (2010). Life-span changes of the human brain white matter: diffusion tensor imaging (DTI) and volumetry. Cereb Cortex, 20(9), 2055–2068. https://doi.org/10.1093/cercor/bhp280

Xu, L., Pearlson, G., & Calhoun, V. D. (2009). Joint source based morphometry identifies linked gray and white matter group differences. Neuroimage, 44(3), 777–789. https://doi.org/10.1016/j.neuroimage.2008.09.051

Yaffe, K., Nasrallah, I., Hoang, T. D., Lauderdale, D. S., Knutson, K. L., Carnethon, M. R., Launer, L. J., Lewis, C. E., & Sidney, S. (2016). Sleep Duration and White Matter Quality in Middle-Aged Adults. Sleep, 39(9), 1743–1747. https://doi.org/10.5665/sleep.6104

Zhang, Y., Brady, M., & Smith, S. M. (2001). Segmentation of brain MR images through a hidden Markov random field model and the expectation-maximization algorithm. IEEE Transactions On Medical Imaging, 20(1), 45–57. https://doi.org/10.1109/42.906424

Zitser, J., Anatürk, M., Zsoldos, E., Mahmood, A., Filippini, N., Suri, S., Leng, Y., Yaffe, K., Singh-Manoux, A., Kivimaki, M., Ebmeier, K., & Sexton, C. (2020). Sleep duration over 28 years, cognition, gray matter volume, and white matter microstructure: a prospective cohort study. Sleep, 43(5). https://doi.org/10.1093/sleep/zsz290

