## Supplementary Table 1 for "The Association Between Poor Sleep and Accelerated Brain Ageing"

**SUPORTING INFORMATION**

**Supplementary Table 1. Complete list of the 110 T1 IDPs that correspond to the volumes of the cortical and subcortical ROIs (GM ROIs) derived from the Harvard-Oxford structural atlases.**

| ID | GM ROI | ID | GM ROI |
|---|---|---|---|
| **1-2** | Bilateral Frontal Pole | 56-57 | Bilateral Cingulate Gyrus (anterior) |
| **3-4** | Bilateral Insula Cortex | 58-59 | Bilateral Cingulate Gyrus (posterior) |
| **5-6** | Bilateral Superior Frontal Gyrus | 60-61 | Bilateral Precuneus |
| **7-8** | Bilateral Middle Frontal Gyrus | 62-63 | Bilateral Cuneal Cortex |
| **9-10** | Bilateral Inferior Frontal Gyrus (pars triangularis) | 64-65 | Bilateral Frontal Orbital Cortex |
| **11-12** | Bilateral Inferior Frontal Gyrus (pars opercularis) | 66-67 | Bilateral Parahippocampal Gyrus (anterior) |
| **13-14** | Bilateral Precentral Gyrus | 68-69 | Bilateral Parahippocampal Gyrus (posterior) |
| **15-16** | Bilateral Temporal Pole | 70-71 | Bilateral Lingual Gyrus |
| **17-18** | Bilateral Superior Temporal Gyrus (anterior) | 72-73 | Bilateral Temporal Fusiform (anterior) |
| **19-20** | Bilateral Superior Temporal Gyrus (posterior) | 74-75 | Bilateral Temporal Fusiform (posterior) |
| **20-21** | Bilateral Middle Temporal Gyrus (anterior) | 76-77 | Bilateral Temporo-occipital Fusiform |
| **22-23** | Bilateral Middle Temporal Gyrus (posterior) | 78-79 | Bilateral Occipital Fusiform Gyrus |
| **24-25** | Bilateral Middle Temporal Gyrus (temporo-occipital) | 80-81 | Bilateral Frontal Operculum |
| **26-27** | Bilateral Inferior Temporal Gyrus (anterior) | 82-83 | Bilateral Central Operculum |
| **28-29** | Bilateral Inferior Temporal Gyrus (posterior) | 84-85 | Bilateral Parietal Operculum |
| **30-31** | Bilateral Inferior Temporal Gyrus (temporo-occipital) | 86-87 | Bilateral Planum Polare |
| **32-33** | Bilateral Postcentral Gyrus | 88-89 | Bilateral Heschl's Gyrus |
| **34-35** | Bilateral Superior Parietal Lobule | 90-91 | Bilateral Planum Temporale |
| **36-37** | Bilateral Supramarginal Gyrus (anterior) | 92-93 | Bilateral Supracalcarine Cortex |
| **38-39** | Bilateral Supramarginal Gyrus (posterior) | 94-95 | Bilateral Occipital Pole |
| **40-41** | Bilateral Angular Gyrus | 96-97 | Bilateral Thalamus |
| **42-43** | Bilateral Lateral Occipital Superior | 98-99 | Bilateral Caudate |
| **44-45** | Bilateral Lateral Occipital Inferior | 100-101 | Bilateral Putamen |
| **46-47** | Bilateral Intracalcarine Cortex | 102-103 | Bilateral Pallidum |
| **48-49** | Bilateral Frontal Medial Cortex | 104-105 | Bilateral Hippocampus |
| **50-51** | Bilateral Juxtapositional Lobule | 106-107 | Bilateral Amygdala |
| **52-53** | Bilateral Subcallosal Cortex | 108-109 | Bilateral Ventral Striatum |
| **54-55** | Bilateral Paracingulate Gyrus | 110 | Brainstem |
